# Extracting Reproducible Components from Electroencephalographic Responses to Transcranial Magnetic Stimulation with Group Task-Related Component Analysis

**DOI:** 10.1101/2025.06.02.657489

**Authors:** Bruno Andry Couto, Matteo Fecchio, Simone Russo, Enrico De Martino, Sara Parmigiani, Simone Sarasso, Thomas Graven-Nielsen, Daniel Ciampi de Andrade, Marcello Massimini, Mario Rosanova, Adenauer Girardi Casali

## Abstract

Transcranial magnetic stimulation combined with electroencephalography (TMS-EEG) is a powerful technique for investigating human cortical circuits. However, characterizing TMS-evoked potentials (TEPs) at the group level typically relies on grand averaging across stimulus repetitions (trials) and subjects – an approach that assumes a level of spatial and temporal consistency that is often lacking in TEPs. Here, we introduce an adaptation of Group Task-Related Component Analysis (gTRCA), a novel multivariate signal decomposition method, to automatically extract TEP components that are maximally reproducible across both trials and subjects. Following the validation of a new permutation-based statistical test for gTRCA using simulated data, the method was applied to two independent TMS-EEG datasets, in which stimulation was targeted to the primary motor cortex (M1) in cohorts of 16 and 22 healthy participants. We found that gTRCA reliably identified TEP components that were reproducible at the group level. Notably, the main gTRCA component captured the key spatial, temporal, and spectral features of motor TEPs, remained robust despite reduced number of stimuli and participants, and was consistent across different recordings. These findings demonstrate that gTRCA affords a more reliable characterization of TEPs at the group level, thereby facilitating the translation of TMS-EEG research into clinical practice.

## Introduction

Combining Transcranial Magnetic Stimulation with electroencephalography (TMS-EEG) enables the non-invasive assessment of cortical excitability and effective connectivity with excellent temporal resolution (Komssi and Kähkönen, 2006; Massimini et al., 2009; Bortoletto et al., 2015; Farzan and Bortoletto, 2022). Over the past two decades, this technique has been instrumental in researching novel biomarkers for a variety of clinical applications (Tremblay et al., 2019). However, its use has been largely confined to the research environment, and its clinical application is still hindered by technical and practical limitations (Julkunen et al., 2022; Bertazzoli et al., 2025). A critical step to bridge the gap between research and clinical applications is to demonstrate that TMS-EEG measures are reliable across cohorts of subjects from independent research groups. Achieving this goal requires simplifying the interpretation of the results of TMS-EEG experiments by reducing the dimensionality of TMS-EEG data, an essential step for highlighting TMS-evoked components most representative of the condition under study while preserving clinically relevant inter-individual differences.

The method commonly employed to reduce the dimensionality of TMS-EEG data is the “grand average”: averaging EEG signals first across trials and then across participants. This practice is widespread in the TMS-EEG community (Belardinelli et al., 2021; Biabani et al., 2019; Chung et al., 2015; Conde et al., 2019; Gordon, Song, et al., 2023; Gordon, Song, et al., 2023; Gordon et al., 2021; Gordon et al., 2018; Guzmán López et al., 2022; Krile et al., 2023; Rocchi et al., 2021), and has also been adopted in recent clinical applications (Bruckmann et al., 2012; Chowdhury et al., 2023; Darmani et al., 2019; Helling et al., 2023; Leodori et al., 2023; Luo et al., 2023; Santoro et al., 2024). However, the grand averaging method implicitly assumes that evoked waveforms recorded at fixed EEG channels are comparable across individuals (Luck, 2014). This assumption may be reasonable for short-latency peripheral evoked potentials, whose intrinsic reproducibility results from the stimulation of anatomical pathways that are invariant across healthy subjects (Chiappa, 1997; Cruccu et al., 2008), but it does not necessarily hold for TMS-evoked potentials (TEPs). The pathways engaged by direct stimulation of cortical circuits with TMS are highly variable across individuals and depend on several parameters – such as stimulation intensity, coil position and orientation– that are difficult to control and standardize at the group level (Belardinelli et al., 2019; Casarotto et al., 2022; Lioumis and Rosanova, 2022; Guidali et al., 2023). Even when state-of-the-art procedures for TMS-EEG acquisition are employed, TEPs often differ in amplitude, waveform, and duration across subjects (Fecchio et al., 2025). In addition, inter-individual variability in head and cortical geometry can cause the same EEG channels to sample different neural sources across subjects (Ozdemir et al., 2021), leading to variable topographic maps at fixed latencies, polarity inversions at fixed channels, and the potential cancellation of meaningful components in the grand average.

In this work, we move beyond the grand average approach and introduce an adaptation of a novel multivariate EEG decomposition technique, called Group Task-Related Component Analysis (gTRCA; Tanaka, 2020), as an alternative dimensionality-reduction method for TMS-EEG data. gTRCA derives subject-specific spatial filters by jointly maximizing temporal correlation across trials and across subjects, thus dispensing with waveform averaging. Because it makes no assumptions about uniform topographies or consistent polarities across subjects, gTRCA should recover group-reproducible TEP components that are more reliable and robust than those obtained by simple grand averages. Here, we test this hypothesis using simulated data and real motor cortex (M1) TEPs from two cohorts of healthy individuals. After validating a new permutation-based statistical test for group-level reproducibility, we examine whether gTRCA components evoked by TMS are (i) reproducible across subjects while retaining known signatures of M1 TEPs in spatial, temporal and frequency domains; (ii) consistent across TMS-EEG datasets obtained by independent research groups; and (iii) robust to variations in the number of trials, participant pool, and sample size.

## Methods

### Group Task-Related Component Analysis

Let us denote an EEG epoch (trial) recorded in a subject *α* (*α* = 1, …, *A*) as the *n × τ* matrix 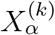, where *k* represents a specific epoch (*k* = 1, …, *K*), *n* the number of EEG channels, and *τ* the epoch length (number of samples). In this section it is assumed that *n, τ*, and *K* are the same for all subjects, but this restriction is enforced only for simplicity of notation and can be relaxed in practice (Tanaka, 2020).

Let us then define *X*_*α*_ *∈* ℝ^*n×Kτ*^ as the *n × Kτ* matrix formed by concatenating all epochs of subject *α* along the temporal dimension and by subsequently normalizing each channel with zero mean and unit standard deviation. In the original description of gTRCA (Tanaka, 2020), *X*_*α*_ represented the raw, continuous EEG, prior to epochs segmentation. Given the heavy artifact burden of TMS-EEG, we inverted this order: we first perform the multiple artifact-removal steps that are commonly used in the preprocessing of TMS-EEG data, including epochs visual screening and Independent Component Analysis, and then concatenate the resulting set of clean epochs to form *X*_*α*_.

From 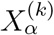, a matrix *S* is defined as follows

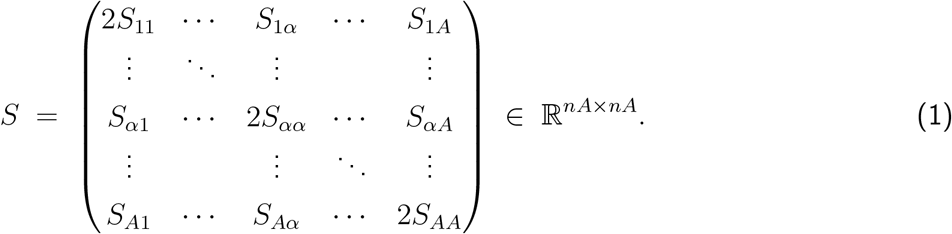

where the off-diagonal elements, *S*_*α,β*_ with *α ≠ β*, represent the inter-subject reproducibility and are defined as the average temporal correlation across all trials of subjects *α* and *β*:

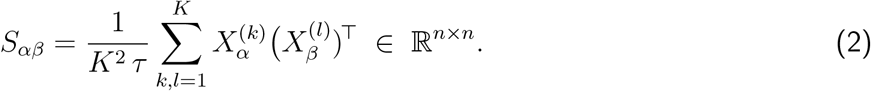

The diagonal elements *S*_*αα*_ represent the reproducibility across trials (within-subject) and are defined as the average correlation across trials for subject *α*:

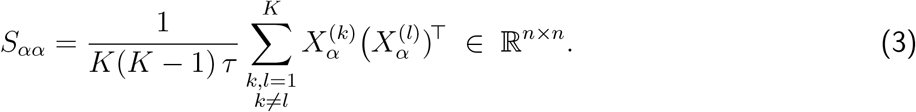

From *X*_*α*_ we can calculate the EEG covariance matrix *Q*_*α*_ of each subject

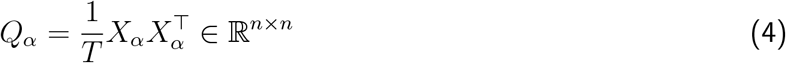

and, following the same block-design of *S*, we can define the full covariance *Q ∈* ℝ ^*nA×nA*^ as the block-diagonal matrix having *Q*_*α*_ (*α* = 1, …, *A*) as its submatrices.

The main goal of gTRCA is to search for temporal components by constructing spatial filters (i.e., by combining EEG channels) so that *S* is maximized when projected onto these filters. Therefore, a set of *A* spatial filters are defined as the following vector *w ∈*ℝ ^*nA×*1^

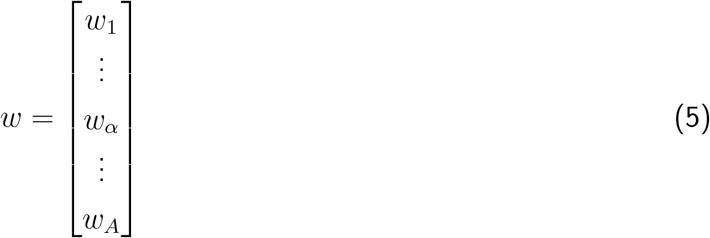

where *w*_*α*_ *∈*ℝ ^*n×*1^ is the spatial filter of subject *α*. We want to find *w* such that *w*^*⊤*^*Sw* is maximum under the constraint that the covariance matrix *Q* is fixed, that is, *w*^*⊤*^*Qw* = 1. This is a Rayleight-Ritz eigenvalue problem, with solution 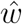 given by

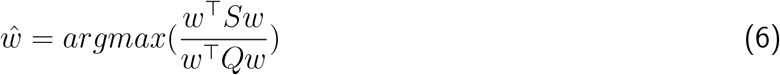

The solutions to this problem can be found by diagonalizing *Q*^*−*1^*S*: the resulting eigenvectors are the projectors *w*, which can then be ordered by their corresponding eigenvalues (*λ*). These eigenvalues represent the strength of *S* when projected onto *w* and are thus measures of overall inter-trial (diagonal terms of *S*) and inter-subject reproducibility (off-diagonal terms of *S*). To account for the scaling of *λ* with the number of subjects (*A*), the normalized eigenvalue *λ*_*A*_ is defined as *λ*_*A*_ = *λ/A*.

Each spatial filter *w* is composed by spatial filters of individual subjects *w*_*α*_, which, in turn, represent a particular gTRCA component 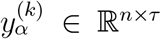, where 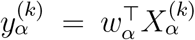. It is important to note that different subjects generally have different spatial filters: in contrast to grand-averages strategies, there is no a priori fixation of EEG channels across subjects. Indeed, each component can be associated to a different scalp-map in each subject, *m*_*α*_ *∈*ℝ ^*n×*1^ where *m*_*α*_ = *Q*_*α*_*w*_*α*_ (Haufe, Dähne, and Nikulin, 2014). A scalp map *m*_*α*_ is a forward operator that projects the corresponding component in the EEG space of a particular subject, thus representing the importance (strength and directions) of the extracted component for each EEG channel of that subject (Haufe, Meinecke, et al., 2014). Different subjects may present topographical distributions for the same gTRCA component with different strengths and directions, including opposite polarities.

The algorithm in Python for extracting the spatial filters, gTRCA components, and scalp maps from epoched data is available at https://github.com/Boutoo/gTRCA. A detailed step-by-step description of the implemented gTRCA class is available in the Supplementary Methods.

### 2.2 GTRCA Statistical Tests

As described in the preceding section, gTRCA finds components by diagonalizing the *Q*^*−*1^*S* matrix. Consequently, the number of components resulting from the method’s application to a given dataset is determined by the rank of *Q*^*−*1^*S* rather than the presence or absence of temporal reproducibility between trials or subjects in the data. EEG recordings that are not reproducible across trials or subjects may have up to *nA* resulting components, but what sets these components apart from those extracted from a dataset that exhibits temporal reproducibility is the magnitude of their corresponding eigenvalues. Therefore, applying an appropriate statistical test to the eigenvalues of the extracted components is a critical step in using multivariate methods like gTRCA with real data. In this section, we introduce two methods for testing the temporal similarity within and between subjects of the components extracted by gTRCA: the first method (trial-based shifting) is grounded on the null hypothesis that there is no temporal locking in the EEG responses among both trials and individuals; the second method (subject-based shifting) employs the null hypothesis that there is no temporal locking between individuals, while allowing for time-locking between trials within each individual.

#### 2.2.1 Method 1: trial-based shifting

In the article introducing gTRCA, Tanaka employed a resampling-based statistical method, previously validated in other studies and designed to test the temporal similarity of components between trials and/or individuals (Tanaka et al., 2013, 2014). Tanaka’s strategy involved randomizing the trials onsets of the continuous EEG recordings for all subjects, thus generating surrogate data not time-locked to the stimuli, and then comparing the eigenvalues of the components in the original data with those in the surrogate data. In the present work, we adapted this method for application to pre-processed and segmented TMS-EEG data using a circular shift of trials: we randomly shifted each trial within a subject along the time dimension in a circular manner, such that samples shifted beyond the end of the time window were introduced at the beginning of the array.

Specifically, we tested the null hypothesis that there is no temporal locking in the data using the following procedure:

1. Each trial for each subject was randomly and independently shifted along the temporal axis. During this process, shifts were random between trials and subjects, but all channels of a given trial and subject were shifted by the same amount. This step results in a surrogate dataset constructed in accordance with the null hypothesis that there is no temporal locking across trials and subjects, while preserving the spatiotemporal structure of EEG signals.

2. gTRCA was then applied to the surrogate dataset, and we extracted the maximum eigenvalue among all components. This procedure is implemented in the method *run surrogate* of the gTRCA class, with parameter mode set to *“trial”* (see the Supplementary Methods for a description of the gTRCA class).

3. The steps above were repeated N times, resulting in a distribution of N maximum eigenvalues under the null hypothesis.

4. Finally, the eigenvalues of the original data were compared to the surrogate distribution and the null hypothesis was rejected whenever the eigenvalues fell within the upper 5% of the distribution.

#### 2.2.2. Method 2: subject-based shifting

Since the null hypothesis of the method described above is the absence of any temporal locking, rejecting this hypothesis implies that a component exhibits significant temporal locking, either across trials or individuals, but not necessarily both. Consequently, a dataset with TMS-evoked potentials lacking any temporal similarity between individuals could result in components that are considered significant by this method solely due to time-locking between trials within each individual.

To address this limitation, we developed a second strategy specifically designed to test temporal reproducibility across individuals. In order to achieve this, we modified the first step of the previous method and shifted all trials of a given subject by the same amount. In this way, by generating surrogate data in which temporal relationships across trials were preserved within each subject but disrupted between subjects, we could directly evaluate the null hypothesis of no temporal similarity between subjects in the original data. This procedure is implemented in the method *run surrogate* of the gTRCA class, with parameter mode set to *“subject”*.

### 2.3 Simulated Data

We applied gTRCA and both statistical methods for identifying reproducible components on simulated EEG datasets. The simulations were generated with MNE-python (Gramfort, 2013) using the template available in MNE (Bekhti et al., 2017). Evoked components were constructed from sine waves within gaussian envelopes that were introduced at the vertices of specific cortical areas. The forward model of the MNE template was used to project the corresponding activation onto 59 channels at the scalp level located according to the 10-20 EEG placement (Bekhti et al., 2017). Trials of 1.2 seconds at 600Hz of sampling frequency were then generated by adding the scalp sinusoids to multivariate Gaussian noise constructed with the estimated EEG covariance of the MNE template and a third-order 60 Hz low-pass i.i.r. filter in both forward and reverse directions. The signals were then re-referenced to the average of all EEG channels. Two distinct sets of simulated data were created:

#### 1. Simulated dataset 1: simulation of reproducibility across trials

19 Hz sinusoids with a duration of 250 ms were introduced in the left postcentral gyrus of the cortical mesh (Figure 1A). EEG signals with 100 trials were generated for a total of 10 simulation runs or “subjects” (see Figure 1B for an example of the evoked potentials in one subject). The polarity of the sinusoids was random between subjects but fixed between trials, and the sinusoids of different individuals were linearly distributed along the duration of the trials to minimize overlaps across subjects (Figure 1C). In this manner, our first simulated dataset ended up composed by signals that satisfy the null hypothesis of the subject-based shifting statistics - as there is no temporal overlap of the evoked components between individuals - but violate the null hypothesis of the trial-based method, since each individual has a component that is temporally invariant across trials.

**Figure 1.**
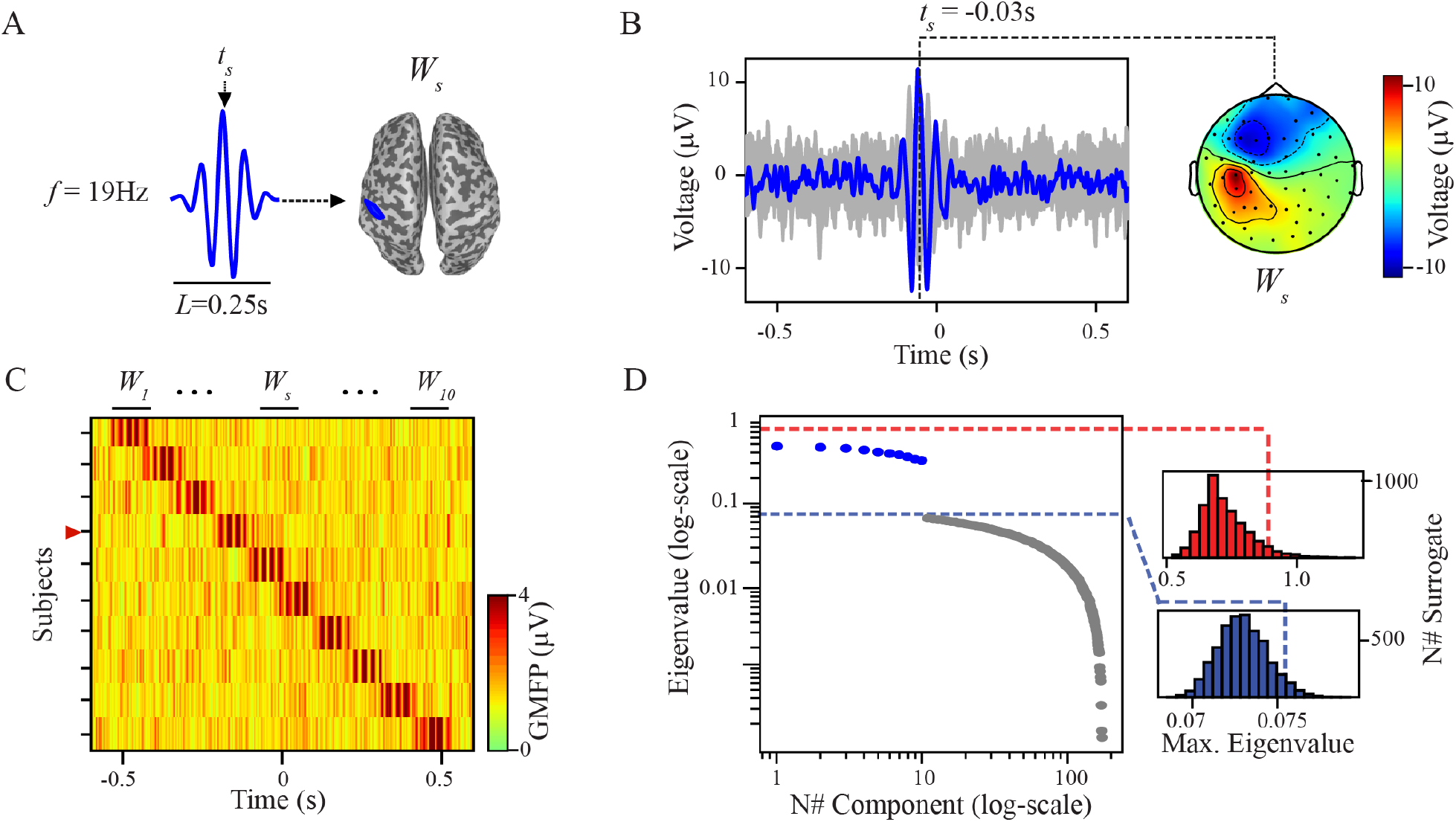
GTRCA and the trial-based shifting test detected all simulated components that were reproducible within but not across subjects. **(A)** Waveforms of the 10 simulated components (*W*_*s*_), each with fixed frequency (*f*) and duration (*L*), but with subject-specific peak latencies *t*_*s*_. Components were localized over the left postcentral gyrus (cortical mesh shown at right) to simulate activity that is temporally invariant across trials of the same subject (*s*) but with minimal temporal overlap across subjects *s* = 1 *−* 10. **(B)** Example of a simulated evoked potential (average of 100 trials) at the scalp level for subject *s* = 4, with *ts* = *−* 0.03*s*. Gray traces represent individual EEG channels, with the channel over the left postcentral gyrus highlighted in blue. The topography at the peak latency of the *W*_*s*_ component is shown on the right. **(C)** Global mean-field power (GMFP), calculated as the voltage root mean square across all channels (Lehmann and Skrandies, 1980, is displayed for all subjects in a heat map; the red marker on the vertical axis indicates the subject shown in (B). **(D)** Eigenvalues of the identified gTRCA components in log scale, along with significance thresholds corresponding to each of the statistical distributions: trial-based shifting in blue and subject-based shifting in red. Eigenvalues of the significant components in the trial-based test are highlighted in blue.

#### 2. Simulated dataset 2: simulation of reproducibility across trials and subjects

The second simulated dataset was constructed by taking the first simulation (described above) and adding a sinusoid, now of 10 Hz, with a duration of 300 ms and over the right precentral gyrus (Figure 2A). This resulted in evoked potentials containing two components for each subject (Figure 2B). The new component was positioned at the same latency in all 10 individuals, such that now, in addition to the components without temporal overlap across subjects, there is a reproducible component that violates the null hypothesis of the subject-based shifting method. Importantly, the polarity of this reproducible component was also randomized between subjects, causing it to be canceled out in the grand-average.

**Figure 2.**
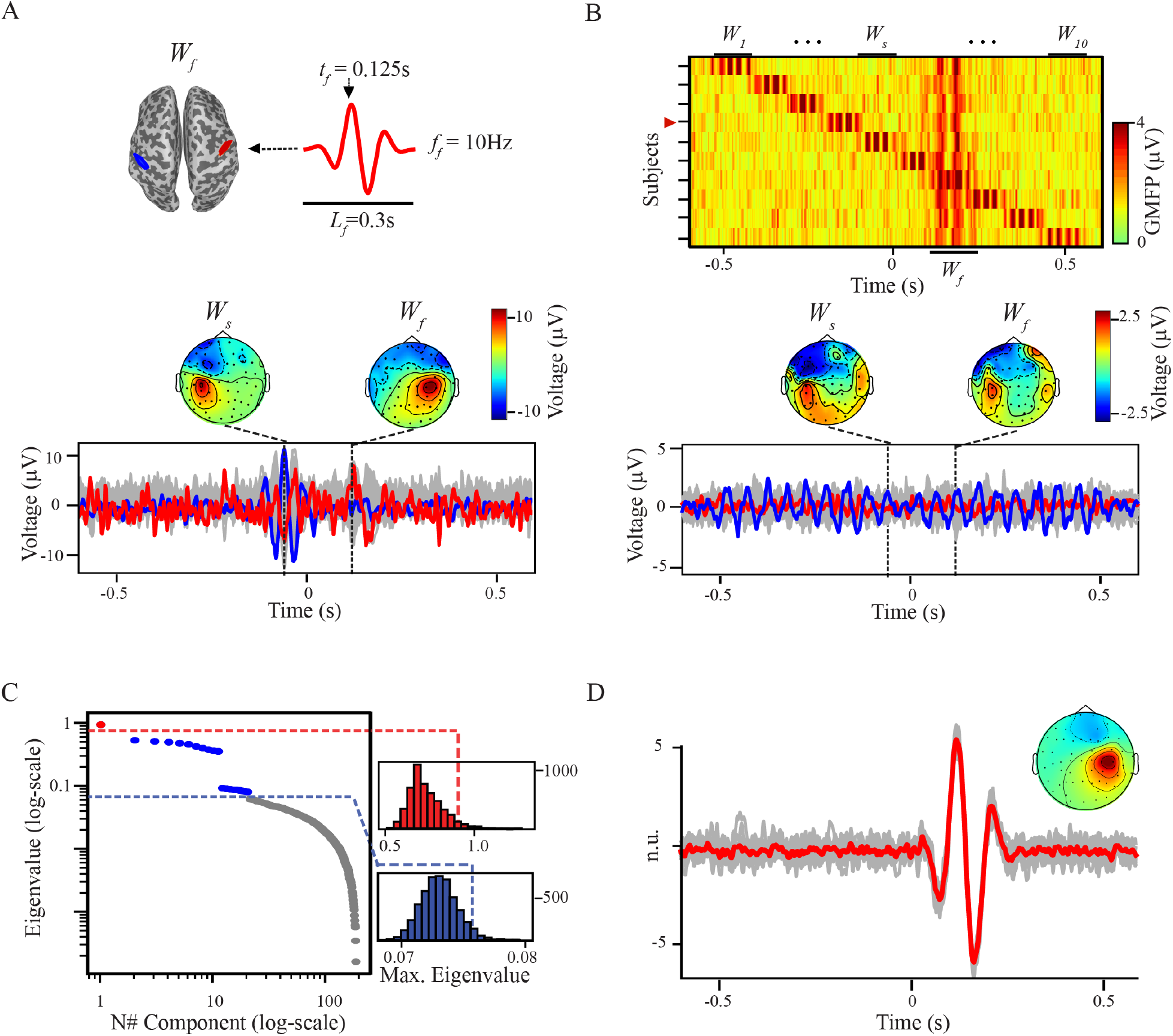
GTRCA and the subject-based shifting test detected the simulated component that was reproducible both within and across subjects. **(A)** Waveform of the added *W*_*f*_ component, reproducible across subjects, introduced at 0.125s over the right precentral gyrus (red region on the cortical mesh) on top of the simulation shown in Figure 1. Below, the evoked potential of the same subject shown in Figure 1B is displayed, now highlighting also a channel over the right postcentral gyrus (in red). Topographies at the peak latencies of both components (*W*_*s*_ as in Figure 2, and *W*_*f*_) are shown above. **(B)** GMFPs for all subjects as a heat map; the red marker on the vertical axis indicates the subject shown in (A). Below, time courses and topographies at the peak latencies of *W*_*s*_ and *W*_*f*_ are shown for the grand-average evoked potential. As the reproducible component has random polarities across subjects, the channel over the right postcentral gyrus (red) exhibits a flat signal after averaging across subjects. **(C)** Eigenvalues of the gTRCA components (log scale) as displayed in Figure 1D, now with one component identified as significant by the subject-based method (in red). **(D)** Waveform and average topographic distribution of the reproducible component identified in (C). Gray traces represent the components for each subject (in normalized units), and the red trace shows the average across subjects.

### 2.4 Real TMS-EEG Data and EEG pre-processing

Two independent TMS-EEG datasets targeting the left primary motor cortex (M1) of healthy individuals were employed in this study. Both protocols followed the same quality-control procedures: real-time monitoring of TMS-evoked potentials with a custom-made graphical user interface (rt-TEP; Casarotto et al., 2022) to ensure a minimum early peak-to-peak amplitude response of 6 *µ*V; auditory masking with the TMS-click sound-masking toolbox (Russo et al., 2022) to suppress auditory artifacts; and continuous electromyography to suppress cortico-spinal contamination. Despite these similarities, the two datasets were recorded and analyzed by independent research groups and acquired with distinct hardware and targeting strategies that introduce meaningful sources of variability. Specifically, they employed different EEG recording systems and stimulator brands as well as different strategies to set the stimulation parameters. Below we briefly describe each protocol. Full methodological details are available in the original publications (De Martino et al., 2023; De Martino et al., 2024; Fecchio et al., 2025).

#### 2.4.1 Milan cohort

TMS-evoked potentials were collected at the University of Milan in 16 healthy individuals (5 female, 2 left-handed, age: 25-51) using a focal figure-of-eight coil (mean/outer winding diameter 50/70 mm, biphasic pulse shape, pulse length 280 *µ*s) connected to a Mobile Stimulator Unit (eXimia TMS Stimulator, Nexstim Ltd) as described in Fecchio et al., 2025. EEG was recorded at 5kHz with a 62-channel cap, plus two channels for electrooculogram, on a TMS-compatible amplifier (BrainAmp DC, Brain Products, GmbH). Ground and reference electrodes were placed on the forehead, and all impedances were kept below 5 kΩ. Individual T1-weighted MRI scans (1.5 T or 3 T) guided coil placement via Nexstim neuronavigation. Stimulation parameters (intensity, coil orientation and position) were determined individually for each participant using rt-TEP (Casarotto et al., 2022). This was done by delivering an initial test pulse at 120V/m over the target area to check for craniofacial muscle activation, and slight adjustments to intensity and coil orientation were made if muscle twitching occurred. Effectiveness in stimulating the cortex was then verified by inspecting the amplitude of early TEP components recorded beneath the TMS coil, and stimulation parameters were slightly adjusted if the peak-to-peak amplitude averaged over 20 trials was below 6*µ*V. To avoid cortico-spinal activation and TMS-evoked muscle contractions, electromyography (EMG) was continuously monitored from six electrodes placed over right-hand muscles (eXimia system, with a sampling rate of 3000 Hz and a cutoff of 500 Hz for low-pass filtering). If motor-evoked potentials (MEPs) were detected, the coil was slightly repositioned or rotated until no TMS-induced motor twitch was recorded. At least 200 pulses were applied to each individual with an inter-stimulus interval randomly jittered between 2000 and 2300 ms (0.4–0.5 Hz). The protocol was approved by the Comitato Etico Milano Area 1 and all participants provided written informed consent. Data was permanently anonymized at the end of recordings and before analyses.

#### 2.4.2 Aalborg cohort

TEPs at Aalborg University were obtained in 22 healthy individuals (11 female, 2 left-handed, age: 20-44) using a Magstim Super Rapid2 Plus1 stimulator with a 70mm figure-of-eight double air film coil. EEG was recorded at 4.8kHz from 63 scalp passive electrodes (EASYCAP GmbH, Etterschlag, Germany) using a g.HIamp EEG amplifier (g.tec-medical engineering GmbH, Schiedlberg, Austria). Electrode impedance was monitored to stay below 5 kΩ. Electrooculogram activity was recorded with two electrodes, positioned laterally to the eyes. An optical-tracking system paired with a navigated brain stimulation system (Brainsight TMS Neuronavigation, Rogue Research Inc., Montréal, Canada) was used to calibrate the participant’s head and TMS coil position. The M1 target was functionally identified by locating each individual’s motor “hot spot” over the left hemisphere’s first dorsal interosseous (FDI) muscle, defined as the site where the largest motor-evoked potential was recorded using EMG electrodes. The resting motor threshold (rMT) was determined as the TMS intensity required to elicit MEPs greater than 50 *µ*V in 5 out of 10 trials measured from the FDI muscle EMG. MEPs were recorded using two electrodes (Ambu Neuroline 720, Ballerup, Denmark) placed parallel to the FDI muscle fibers, with a reference electrode positioned on the ulnar styloid process. The stimulation intensity was set to 90% of the rMT to avoid contamination from sensory feedback. At least 160 pulses were applied to each individual with an inter-stimulus interval randomly jittered between 2600 and 3400 ms (0.3–0.4 Hz) The study was approved by the Institution’s Ethics Review Board (Den Videnskabsetiske Komité for Region Nordjylland: N-20220018) as part of the baseline assessment of a research protocol in healthy participants (De Martino et al., 2023; De Martino et al., 2024). Data was permanently anonymized at the end of the recordings and before analyses.

#### 2.4.3 Data preprocessing

Preprocessing was performed independently by each research group, but using the same analysis pipeline. EEG signals were preprocessed in Matlab (The MathWorks, Inc., Natick, MA, United States) versions R2016b (Milan) and R2019b (Aalborg). After visual inspection to reject epochs and channels contaminated by artifacts, between 164 and 273 (218 ± 30) epochs were retained in the Milan data and between 110 and 181 (158 ± 16) in the Aalborg data. Overall, no more than four channels were removed for each session in both datasets. Artifacts associated with the magnetic pulse were removed by replacing the interval between −2ms and 8ms with the immediately preceding interval, followed by a 5th-order moving-average filter applied between 6 and 10ms. Signals were then filtered with a high-pass filter at 1Hz, segmented into 1.6-second windows centered on the stimulation time, and referenced to the average of all good channels (average reference). Independent Component Analysis (ICA) was used to remove artifacts associated with eye movements, spontaneous muscle activity, and residual magnetic artifacts. At the end of the process, a 3rd order Butterworth low-pass filter at 45 Hz was applied in both directions, followed by downsampling to 500Hz (Milan) or 400Hz (Aalborg), data segmentation between –0.6s and 0.6s, baseline correction and channels interpolation (EEGLAB spherical interpolation, Delorme and Makeig, 2004).

Time-frequency decomposition of TMS-EEG signals were calculated by using Morlet wavelets with 3.5 cycles between 10 and 50 Hz as described in previous work (Ferrarelli et al., 2012; Rosanova et al., 2009). Characterization of the responses at the group level was performed by concatenating all trials from all subjects for each individual channel and subsequently extracting grand-average event-related spectral perturbation (ERSP) maps as the average ratio between post-stimulus and pre-stimulus spectral power. The same procedure was employed to calculate the ERSP maps of the gTRCA components, but in this case after concatenating all trials from all subjects for each individual component.

### 2.5 Statistical comparison

#### 2.5.1 Quantification of signal similarity

The similarity of a group of signals was quantified by computing the mean pairwise Pearson correlation and a non-parametric confidence interval. Pearson coefficients *r* were variance-stabilised with Fisher’s transform, *z* = atanh(*r*), and the mean correlation point estimate was calculated as the hyperbolic tangent of the mean *z* value. Uncertainty was evaluated by resampling the vector of *z* values with replacement (5000 iterations). In each iteration the resampled *z* values were averaged, converted back to correlation values and stored. The empirical 2.5th and 97.5th percentiles of the resulting distribution provided the two-sided 95% confidence interval. Results are presented as mean correlation (r) and confidence interval [95% CI].

This procedure was applied in three modalities: (i) Channel-wise: to evaluate how accurately the grand-average TEP represented each participant’s evoked response; (ii) Pairwise across subjects: to evaluate the overall similarity of waveforms and spatial maps of components or evoked potentials within a cohort; (iii) Component-wise: to evaluate how accurately average gTRCA components represented the corresponding components in each participant, and to assess the robustness of average waveforms and spatial maps to changes in the number of trials and participant pool.

The robustness of gTRCA components to changes in trial count was tested by recalculating gTRCA after randomly downsampling the data to subsets of *N* trials (*N* = 10–150, in 10-trial increments). At every *N*, 100 independent random subsets were drawn. To assess the effect of sample size, we recomputed gTRCA using all trials but on smaller subsets of subjects (*A* = 1–15 for Milan, *A* = 1–21 for Aalborg). When the number of possible subsets was fewer than 600 (Milan, *A <* 4 and *A >* 12; Aalborg, *A <* 3 and *A >* 19) we tested every combination of subjects, otherwise we drew 600 random subsamples of participants. In all cases, we computed the temporal and spatial correlations between the average components obtained from the reduced subsets and those obtained from the full dataset.

#### 2.5.2 Permutation-based comparison of Milan and Aalborg time series

To test for between-cohort differences we used a non-parametric permutation test that randomises subjects. Under the null hypothesis that the two datasets differ only in sample size, participants were randomly reassigned (500 permutations) to surrogate “Milan” and “Aalborg” groups while preserving the original group size. For every permutation we recomputed (i) the grand-average TMS-evoked response at the channel located beneath the stimulator and (ii) the first gTRCA component. We then calculated the difference between the two surrogate groups for each signal of interest. The empirical two-tailed p-value for the true group difference was obtained by ranking the observed difference within the permutation distribution. Significance was set at each time sample at the 0.05 level after false-discovery-rate correction. For this comparison, EEG recordings from the Aalborg dataset were time-interpolated to match the sampling frequency of the Milan recordings.

## 3 Results

### 3.1 GTRCA identified reproducible components in simulated evoked potentials

We evaluated gTRCA in combination with the two permutation tests - trial-based and subject-based shifting - using two simulated TMS-EEG datasets.

In the first dataset a sinusoidal evoked component *W*_*s*_ was inserted into the left post-central gyrus (Figures 1A–B). *W*_*s*_ waveforms had identical frequency, amplitude and duration across all ten subjects, but different peak latencies, resulting in minimal temporal overlap between subjects (Figure 1C). Running gTRCA on these data yielded 176 components. As expected, the trial-based shifting test identified the first ten components as significant (*p <* 0.05), given that each one of the ten subjects contained a time-locked component reproducible across trials. However, trial-level significance does not imply group-level reproducibility, and the subject-based shifting test-which preserves temporal locking within but not across subjects - found no significant components (Figure 1D).

In the second simulated dataset, a component *W*_*f*_, reproducible across subjects, peaking at 125 ms and located under the right precentral gyrus (Figure 2A, top), was added to the first simulated dataset. As a result, the evoked potentials of each subject consisted of two time-locked components (Figure 2B, top): one locked across trials within each subject but not across subjects, and another time-locked across both trials and subjects. Importantly, the grand-average activity (Figure 2B) failed to reveal the reproducible component, as its polarity was randomized across subjects. However, gTRCA successfully accounted for the opposite polarities. While the trial-based statistical method identified several significant components, the subject-based test identified exactly one significant component (Figure 2C), which had the same latency, waveform, and topography of *W*_*f*_.

### 3.2 Grand averaging failed to capture consistent TEP patterns across datasets

TEPs were highly heterogeneous across subjects within each cohort of participants. The global mean-field power (GMFP; Lehmann and Skrandies, 1980) revealed substantial differences in both amplitude and duration of M1 TEPs among participants in the Milan and Aalborg datasets (Figure 3A). Similar heterogeneity emerged in the spatial domain at fixed latencies: scalp maps at the GMFP peaks (Milan: 50, 95, 120 ms; Aalborg: 48, 110, 170 ms), showed marked between-subject variability, including frequent polarity inversions (Figure 3B). We quantified how similar individual TEPs were to one another by computing, for each channel, the Pearson correlation between all possible pairs of participants and then averaging these values across channels. Pairwise average correlations were low overall and significantly lower in the Aalborg cohort (Milan: *r* = 0.275 [95% CI: 0.242 *−* 0.308]; Aalborg: *r* = 0.172 [0.150 *−* 0.194]).

**Figure 3.**
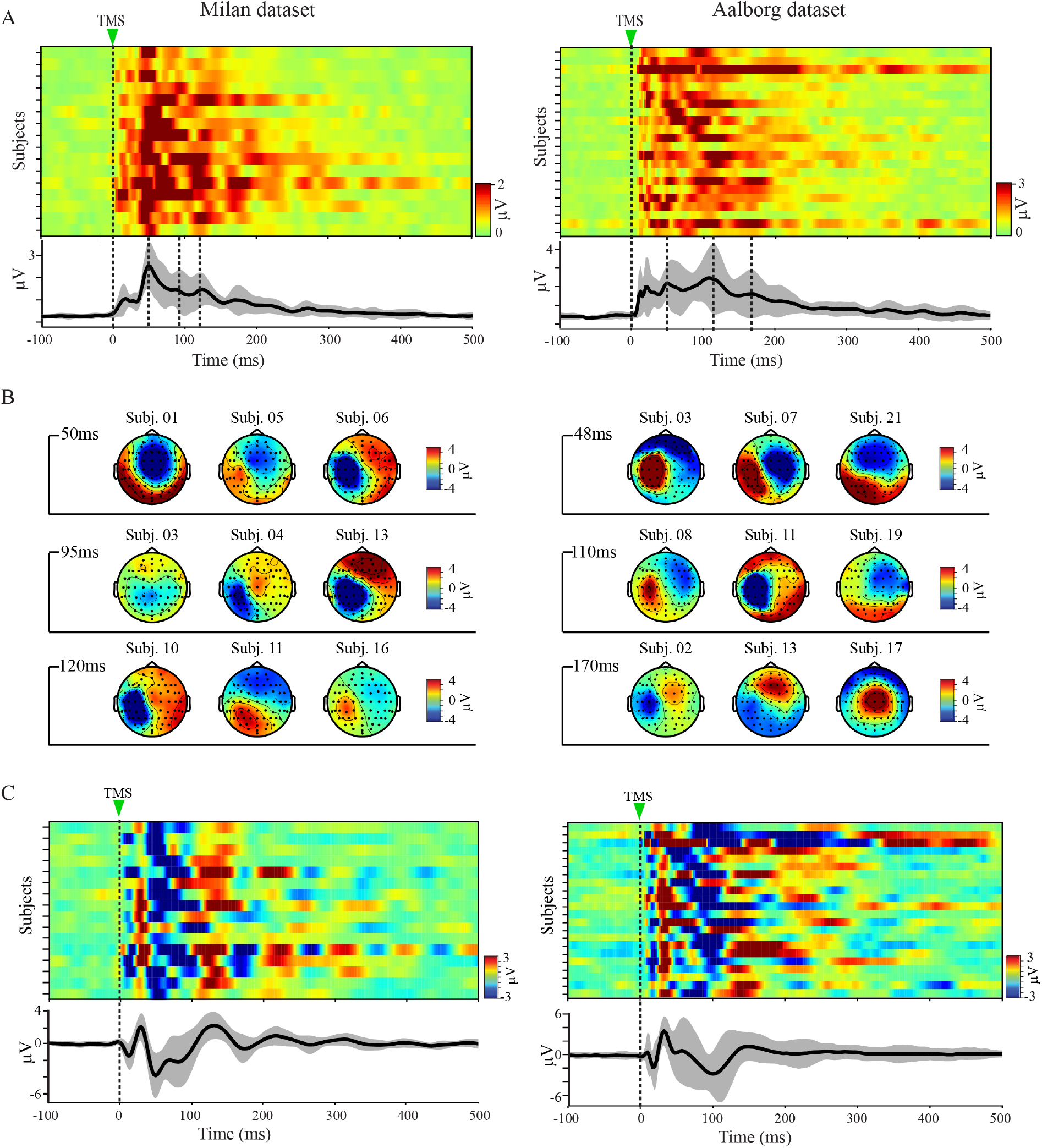
TMS of M1 elicited diverse evoked responses across individuals. **(A)** Global cortical responses to TMS for every participant in the Milan (left) and Aalborg (right) datasets, quantified as global mean field power (GMFP) and shown as heat maps. The black curve and gray shading below depicts the cohort mean ± standard deviation. Dashed vertical lines mark the TMS pulse (0 ms) and selected GMFP peaks. **(B)** TMS-evoked scalp voltage maps at the latencies of the GMFP peaks for representative subjects from the Milan (left) and Aalborg (right) cohorts. **(C)** Heat maps displaying the evoked voltage at channel C1, located directly beneath the coil, for each subject in the Milan (left) and Aalborg (right) cohorts. The black curve and gray shading below show the grand-average waveform ± standard deviation for each cohort.

These global effects, averaged across all channels, were also evident locally at the stimulation site. Polarity, amplitude, duration, and waveform shape of evoked responses at the EEG channel C1, positioned directly beneath the coil, all fluctuated considerably across subjects (Figure 3C). The resulting pairwise correlations for channel C1 were modest on both datasets and significantly lower in the Aalborg cohort (Milan: *r* = 0.446, [0.399 *−* 0.491]; Aalborg: *r* = 0.244 [0.197 *−* 0.291]). Overall, both cohorts presented considerable inter-subject variability, with the Aalborg dataset exhibiting significantly less homogeneity than the Milan cohort.

Time courses and scalp maps of the grand-average TEPs for Milan (Figure 4A) and Aalborg (Figure 4B) datasets exhibited canonical features of M1 TEPs: early, focal peaks over the stimulated area followed by broader components extending up to 400 ms. Yet, given the large inter-subject variability of evoked responses illustrated in Figure 3, these grand-average waveforms cannot be assumed to represent individual participants accurately.

**Figure 4.**
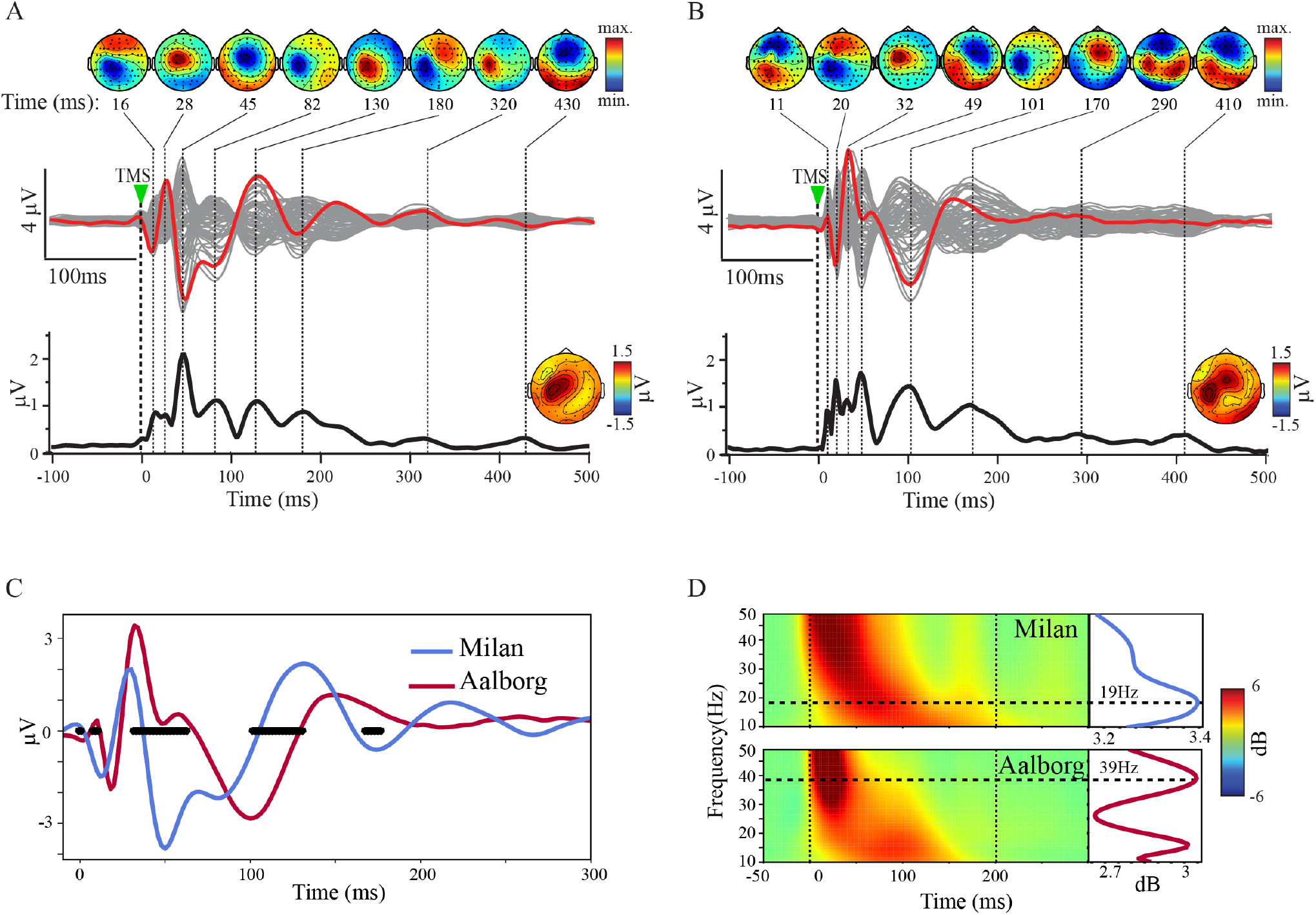
Grand-average TEP evoked by M1 stimulation showed inconsistent waveforms and topographical distributions across different datasets. **(A-B)** Grand averages of the Milan (A) and Aalborg (B) cohorts are displayed in butterfly plots. Individual EEG channels are colored in gray, with the red curves highlighting the electrode under the stimulator (C1). Below, the black curves show the corresponding GMFPs. Topographic maps depict the average post-stimulus mean field power spatial distribution. Above, scalp voltage maps of the grand-average responses are shown for the latencies of the GMFP peaks. **(C-D)** Direct comparisons between the local responses evoked by TMS beneath the coil (channel C1) in time (C) and frequency (D) domains. Evoked responses in the time domain are shown for the Milan (blue) and Aalborg (dark red) cohorts, together with latencies of significant differences in the permutation statistics across groups (black horizontal dots, *p <* 0.05 after FDR correction). Spectral profiles of the same channel depict different time-frequency maps (D, left) and average spectra (D, right) of the first 200ms (Rosanova et al., 2009), peaking at different frequencies (19 Hz for the Milan data, blue curve; 39 Hz for the Aalborg data, dark red curve).

We assessed how well the grand average method captured the group of TEPs by calculating Pearson correlation coefficient between the grand-average and individual responses for each dataset. Globally, the average channel-wise correlation was *r* = 0.577 [0.494 *−* 0.646] for the Milan cohort and *r* = 0.449 [0.399 *−* 0.392] for the Aalborg cohort. Locally, at the channel beneath the TMS coil, mean correlation between the grand-average TEP and individual evoked responses was *r* = 0.688 [0.606 *−* 0.748] for Milan and *r* = 0.494 [0.382 *−* 0.583] for Aalborg. In general, the more heterogeneous Aalborg cohort had a significantly less representative grand-average TEP.

As a result of these differences, marked disparities emerged when grand-average responses were compared across datasets. The evoked signals beneath the coil and the corresponding GMFPs peaked at different latencies and exhibited distinct topographic distributions, as shown in Figures 4A and 4B. When directly compared, the two grand-average TEPs at channel C1 were found to be substantially different in both time and frequency domains (Figure 4C): within the first 200 ms, permutation statistics identified significant differences between the waveforms, and cross-datasets correlations were low (Pearson *r* = 0.23 for the time series; *r* = *−*0.11 for the spectra).

### 3.3 GTRCA identified reproducible components reflecting key features of motor TEPs

GTRCA and the two proposed statistical methods were tested on the Milan dataset, resulting in 224 components (Figure 5B), of which 19 were statistically significant in the trial-based approach (eigenvalues *λ ≥* 0.22). Notably, three of these components were also significant in the subject-based statistics (*λ* = 2.64, *p <* 0.0002; 1.94, *p <* 0.0002; 1.58, *p <* 0.004). The first two components exhibited average waveforms that were highly correlated with the corresponding individual subject waveforms (Figure 5C). The average Pearson correlation was *r* = 0.817 [0.768 *−* 0.861] for the first component and *r* = 0.792 [0.753*−*0.828] for the second. Both values were significantly higher than the average correlation between the grand-average response and individual TEPs beneath the TMS coil. The third component presented an average correlation of *r* = 0.731 [0.640 *−* 0.807].

**Figure 5.**
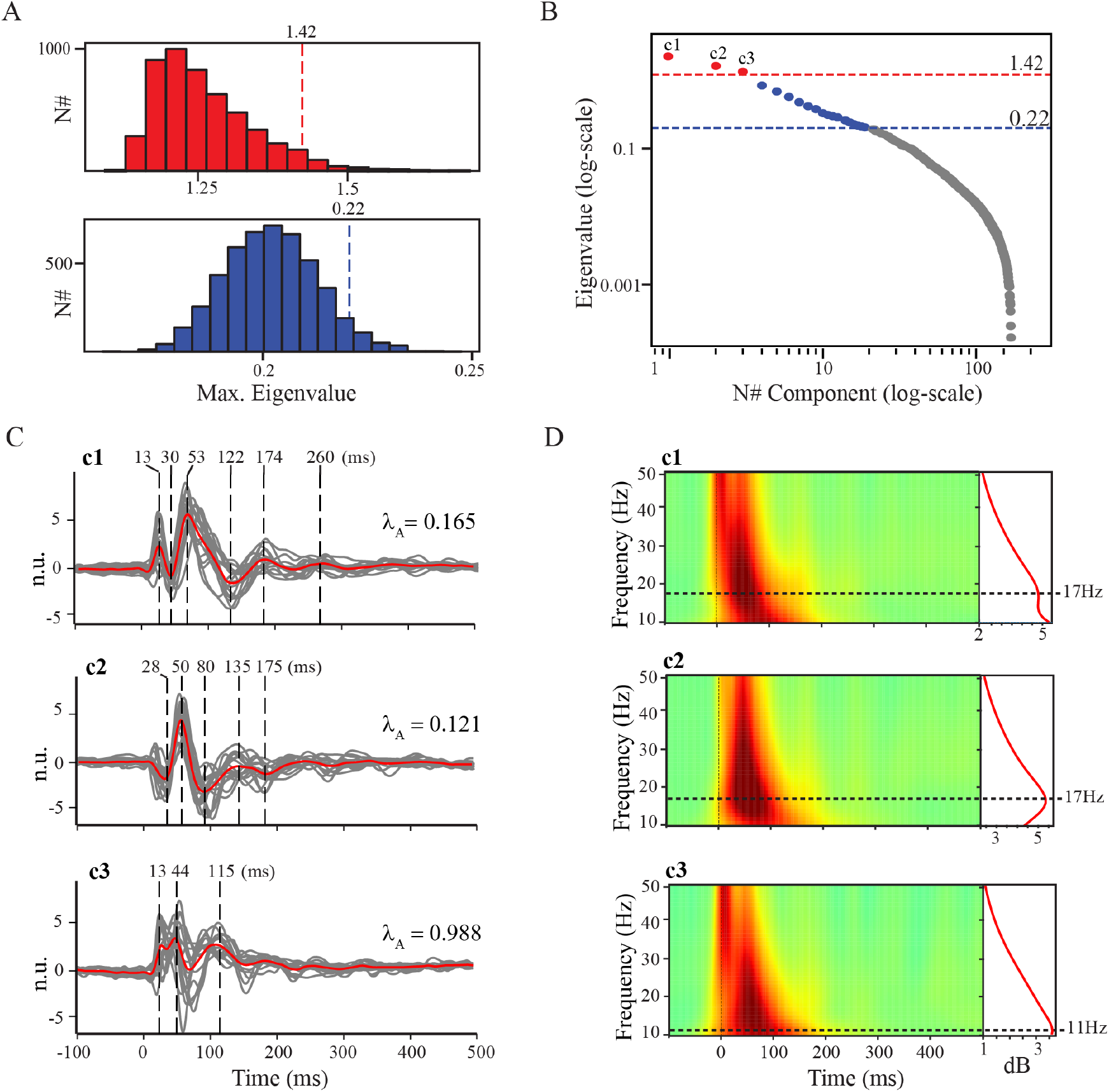
GTRCA extracted TMS-evoked components in the Milan cohort that were reproducible at the group level. **(A)** Statistical null-distributions of gTRCA eigenvalues for each method: trial-based shifting (blue) and subject-based shifting (red). The dashed lines indicate the 95% quantiles, with the corresponding values of the threshold exhibited on top. Both distributions were constructed with 5,000 surrogates (see Supplementary Figure 1 for results with reduced number of surrogates). **(B)** Eigenvalues (in logarithmic scale) obtained from the application of gTRCA. Components that were found not significant in both tests are colored in gray, those that were significant only in the trial-based shifting are colored in blue, and those that were significant in both tests (trial and subject-based shifting) are colored in red and marked as c1, c2 and c3. Vertical dashed lines exhibit the thresholds for the trial-based (blue) and subject-based (red) shifting methods. **(C)** Time series of the three reproducible components, c1, c2, c3, corresponding to the red circles in (B), oriented such that the peak of the early response (0-100ms) has positive polarity, along with their respective normalized eigenvalues (*λ*_*A*_). The time series for each individual is presented in gray, and the average across subjects is shown in red, with main peaks marked by dashed vertical lines. **(D)** ERSP heat maps for each significant component. Red curves on the right display the corresponding average spectra across the first 200ms, and the locations for the peaks of each spectrum are highlighted by the dashed horizontal lines.

Individual waveforms of the first two components were also highly correlated across subjects. Average pairwise Pearson correlation coefficients were 0.656 [0.626 *−* 0.686], 0.630 [0.599 *−* 0.659] and 0.456 [0.394 *−* 0.515], for the first, second and third components, respectively (see Supplementary Figure 2A for full distributions).

The first component displayed prominent peaks at latencies P13, N30, P53, N122, P174, and P260. The second peaked at N28, P50, N80, P135 and N175, while the third peaked at P13, P44, and P115. Time-frequency maps (Figure 5D) revealed an early broadband response followed by activity lasting around 200 ms in all components, and concentrated in the beta-range (peak at 17 Hz) for the first two components and in the alpha-range (peak at 11 Hz) for the third component.

In the spatial domain, all significant components showed an average topographic distribution lateralized over the left hemisphere and centered above the stimulated area (Figure 6). The first two components showed high spatial consistency across subjects, with average pairwise correlation coefficients of 0.638 [0.578 *−* 0.693] and 0.645 [0.586 *−* 0.697], respectively (full distributions in Supplementary Figure 2B). The third component had significantly lower spatial reproducibility, with an average pairwise correlation of 0.359 [0.288 *−* 0.425]. Importantly, although largely reproducible across subjects, the individual spatial maps also enabled identification of subjects who deviated from the group pattern. For instance, participants 1, 3 and 13 showed a first component with a more medial and less lateralized topographical distribution compared to the others.

**Figure 6.**
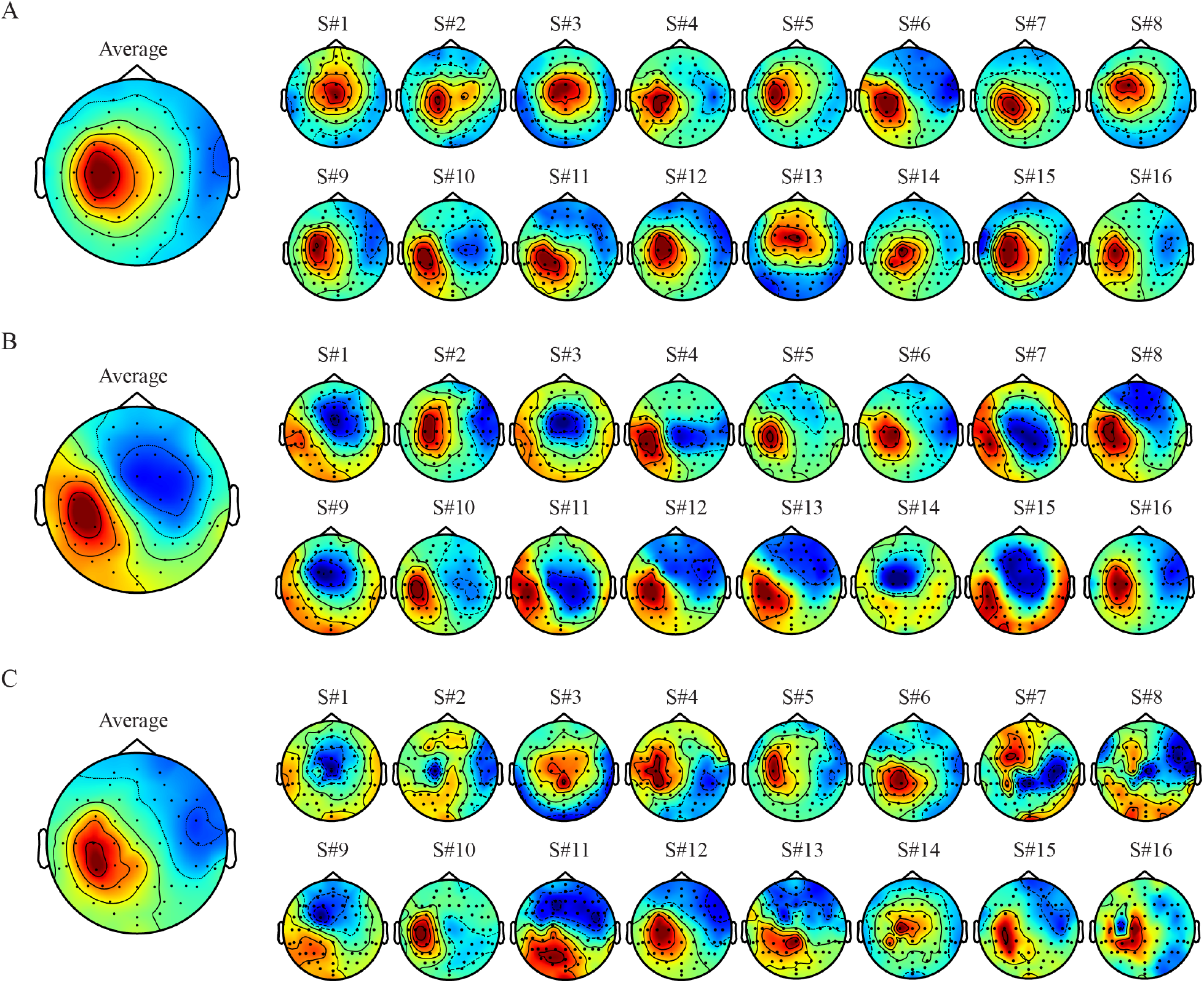
Significant TEP components extracted by gTRCA in the Milan cohort demonstrate spatial maps located over the stimulated region and largely reproducible across subjects. **(A-C)** Scalp maps for each significant component presented in Figure 5 (A: c1; B: c2; C: c3). Averages across all subjects are shown in the left topographical plots, with the corresponding individual topographical maps displayed to the right.

### 3.4 Principal gTRCA components were consistent across datasets

When applied to the Aalborg dataset, gTRCA resulted in 305 components, of which 19 were statistically significant in the trial-based approach (eigenvalues *λ ≥* 0.31). Consistent with the findings from the Milan cohort, three of these components were identified as significantly reproducible based on the subject-based test (*λ* = 3.76, *p <* 0.0002; 2.83, *p <* 0.0002; 2.10, *p* = 0.04 - see also Supplementary Figure 3A-B for eigenvalue distributions). The corresponding normalized eigenvalues were comparable to those observed in the Milan cohort (Figure 5C): *λ*_*A*_ = 0.171, 0.128 and 0.095 for the first, second and third components, respectively.

In the time domain (Figure 7A), the first component peaked at latencies P12, N24, P58, N110, P165, and P247; the second at P13, N30, P90 and N150; and the third at P13, N42, P65, N114, and P165. Average waveforms were strongly correlated with the corresponding individual waveforms for each subject: mean Pearson correlation were 0.806 [0.768 *−* 0.837], 0.798 [0.771 *−* 0.824] and 0.719 [0.684 *−* 0.751] for the first, second and third components, respectively. These values were similar to those obtained in the Milan cohort and significantly higher than the average correlation between the grand-average TEP and single-subject responses beneath the TMS coil. Pairwise inter-subject correlations within the Aalborg dataset were likewise comparable to those observed in the Milan cohort (first, second and third components: 0.650 [0.627 *−* 0.672], 0.656 [0.635 *−* 0.676] and 0.502 [0.475 *−* 0.529]).

**Figure 7.**
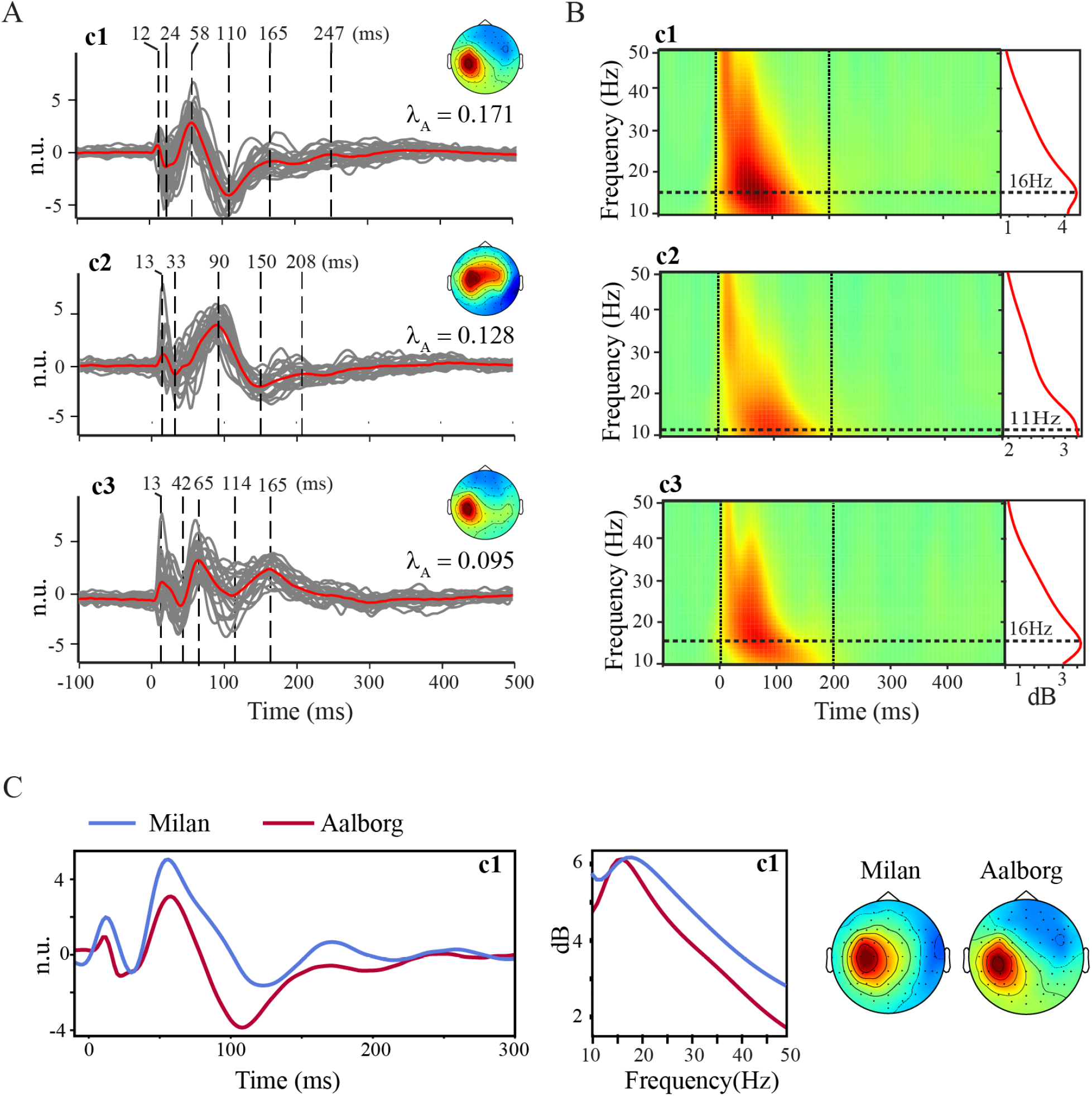
GTRCA identified reproducible components in the Aalborg TMS-EEG dataset that have spatial, temporal and spectral features similar to those in the Milan data. **(A-B)** Time series (A) and ERSP maps (B) of the significant components identified by gTRCA in the Aalborg dataset are shown as in Figure 5 for the Milan cohort (see legend of Figure 5 for further details). Average spatial maps for each component are displayed to the right of panel A. (**(C)** Direct comparison between the first gTRCA component (c1) of the two datasets in time (left - Milan: blue; Aalborg: dark red), frequency (middle - average spectra of the early responses) and spatial (right) domains.

Average scalp maps (right panels in Figure 7A, see also Supplementary Figure 3C) were all lateralized over the left hemisphere and centered above the stimulated area, replicating the Milan pattern (Figure 6). Mean pairwise spatial correlations were 0.651 [0.614 *−* 0.685], 0.471 [0.418 *−* 0.522] and 0.287 [0.223*−*0.349], for the first, second and third component, respectively (full distributions in Supplementary Figure 4).

Time-frequency maps (Figure 7B) also paralleled the Milan results (Figure 5D): the first and third components displayed dominant activity in the beta-range (peak at 16 Hz), whereas the second component showed a peak in the alpha-range (11 Hz).

Overall, despite the inter-subject variability highlighted in Figure 3 and the differences between the two grand-average TEPs (Figure 4), gTRCA was able to extract a common reproducible pattern in both cohorts: the most reproducible gTRCA component (c1) extracted from the Aalborg cohort closely matched its Milan counterpart across spatial, temporal and frequency domains (Figure 7C). The two waveforms correlated strongly within the first 300 ms (*r* = 0.77), and their average power spectra were nearly identical (*r* = 0.97). Permutation statistics detected no significant differences between the two time series.

### 3.5 Principal gTRCA components were robust to variations in trial count and participant pool

We evaluated whether the waveforms and scalp maps of the reproducible gTRCA components depended on the number of trials and the specific composition of participant pools.

Figure 8 shows the mean temporal and spatial correlations between average components derived from a reduced number of trials and those obtained with all trials. In the Milan cohort, the first and second components reached the stability threshold - defined as average spatial and temporal correlation exceeding 0.95 - with 40 and 80 trials, respectively. The third component, however, never achieved an average spatial correlation above 0.95. In the Aalborg cohort, the larger sample size allowed the first two components to stabilise with only 30 trials, while the third required at least 110 trials. Supplementary Figure 5 illustrates how components eigenvalues varied as a function of the number of trials for each cohort.

**Figure 8.**
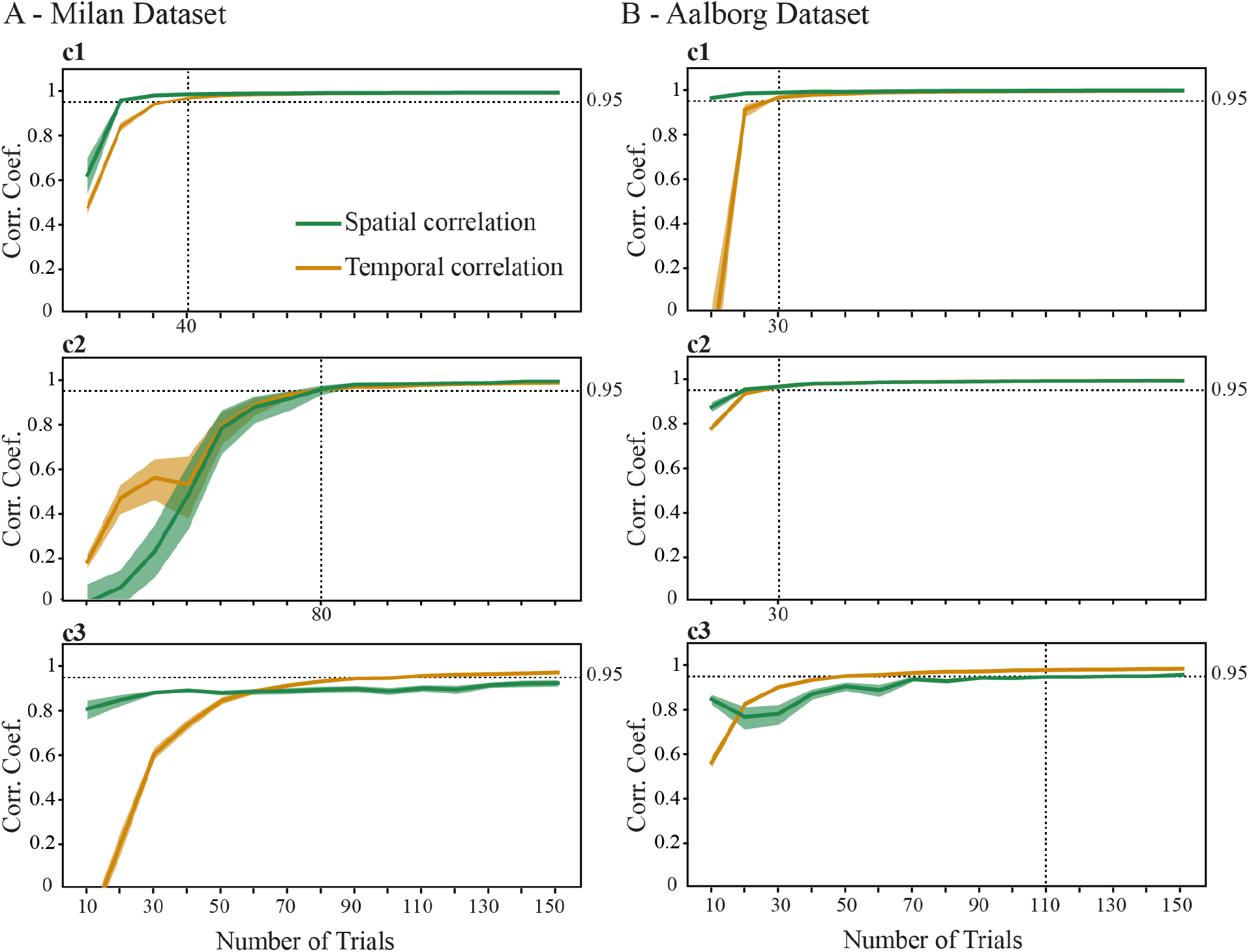
Stability of group-level reproducible components evoked by M1 stimulation as a function of reduced number of trials. **(A-B)** Mean spatial (green) and temporal (yellow) correlation coefficients between full-data average components and the corresponding components recomputed with a reduced number of trials (x-axis). Results are shown separately for the Milan (A) and the Aalborg (B) cohorts. Rows correspond to the first (c1, top), second (c2, middle), and third (c3, bottom) components, respectively. Horizontal dotted lines indicate the minimum number of trials required for the components to achieve waveform and topographic stability, defined as an average temporal and spatial correlation coefficient of at least 0.95. Shaded bands denote the 95% confidence interval.

When the size and composition of participant pools were varied, the 0.95 stability threshold was reached in the Milan dataset with just 8 participants for the first component, while the second and third components required 14 subjects (Figure 9A). In contrast, the significant components of the Aalborg cohort, which had a lower trial count, reached the same level of stability with 13, 17, and 21 participants, respectively.

**Figure 9.**
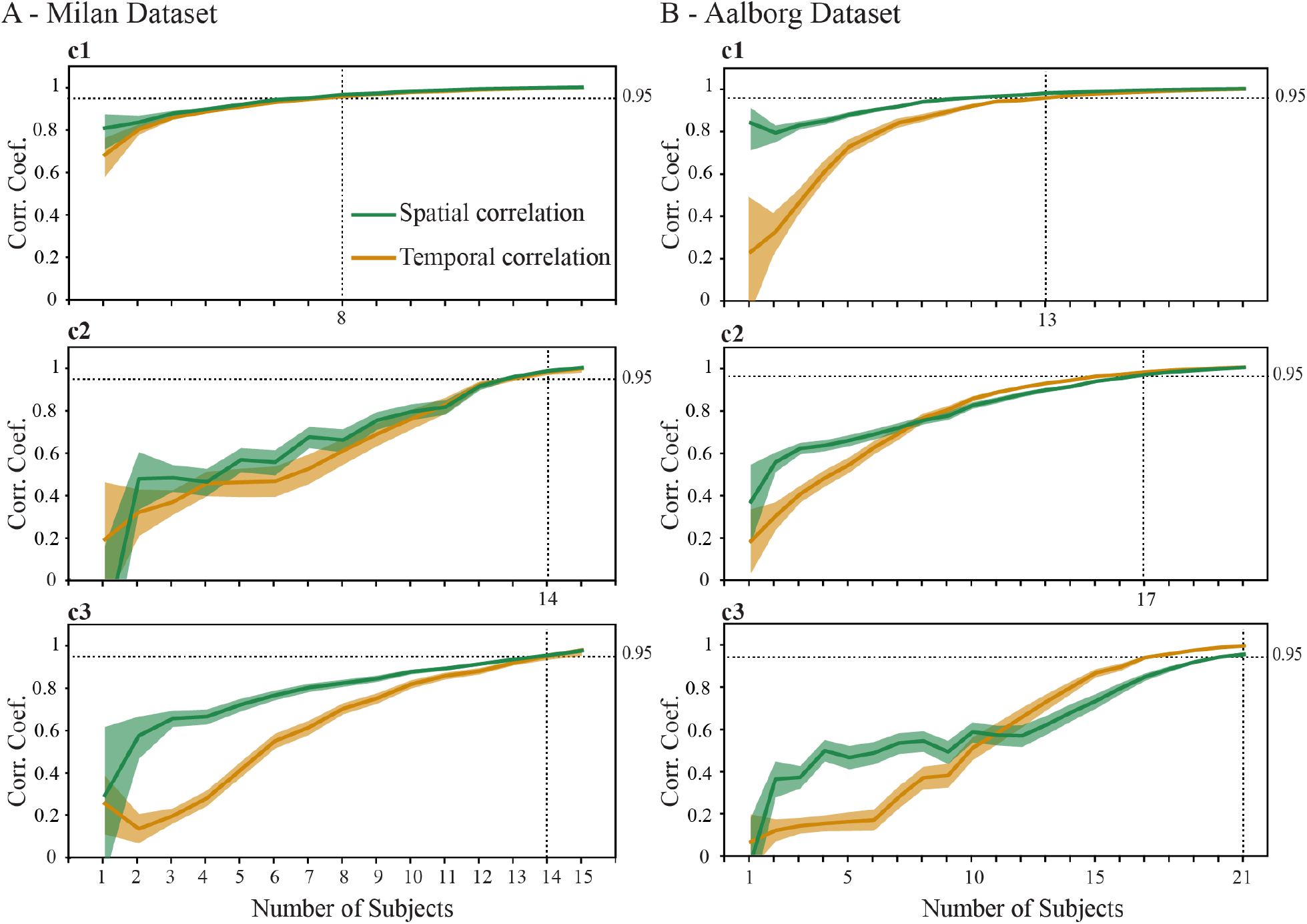
Stability of group-level reproducible components evoked by M1 stimulation as a function of reduced number of subjects. **(A-B)** For the Milan (A) and the Aalborg (B) cohorts, we repeatedly drew up to 600 random sub-samples of n subjects (x-axis) and recomputed the first three gTRCA components. Solid lines show the mean Pearson correlation between each resampled average component and the corresponding component obtained from the full cohort. Spatial correlations are plotted in green, temporal correlations in yellow. Shaded bands denote the 95% confidence interval. Rows correspond to the first (c1, top), second (c2, middle), and third (c3, bottom) components, respectively. Horizontal dotted lines indicate the minimum number of subjects required for the components to achieve waveform and topographic stability, defined as an average temporal and spatial correlation coefficient of at least 0.95.

Finally, we asked whether any single participant could drive components’ significance at the group level. A leave-one-out procedure was run in which gTRCA was refitted N times, each time omitting one subject and regenerating the subject-based null distribution (500 surrogates per iteration, which proved sufficient in Supplementary Figure 1). In every iteration the first and second components remained consistently significant in both datasets (*p <* 0.002), confirming their robustness. The third component was more sensitive, losing significance in 2 of 16 runs for the Milan cohort and in 11 of 22 runs for the Aalborg cohort.

## 4 Discussion

In this work, we showed that gTRCA, coupled with a new subject-based permutation test, reliably isolated motor-cortex TMS-evoked components that were reproducible at the group level, despite large inter-subject variability that undermined conventional grand averaging. The main gTRCA component successfully generalized across two independent cohorts and remained robust with small trial counts and reduced number of participants, demonstrating that the method is a powerful dimensionality reduction technique to deliver cross-dataset biomarkers from TMS-EEG recordings.

### 4.1 GTRCA as a method for TMS-EEG dimensionality reduction

A hallmark of a complex adaptive system such as the brain is the contrast between the large number of its constituent elements and the low effective dimensionality of the system’s emergent large-scale dynamics (Chialvo, 2010; Gotts et al., 2020; Shine et al., 2019; Thibeault et al., 2024). Proper characterization of the low-rank brain dynamics requires using dimensionality reduction techniques that capture the brain activity patterns effectively supporting brain functions on a large scale (Cunningham and Yu, 2014). In this work, we tackled the problem of reducing the dimension of brain signals obtained through the combination of TMS and EEG, a technique that is increasingly being used to explore brain dynamics in search of biomarkers for different neurological and psychiatric conditions (Farzan, 2024; Tremblay et al., 2019).

Most TMS-EEG studies reduce data dimensionality at the group level by grand averaging signals across trials and subjects. Although this practice has yielded valuable insights into the characterization and potential applications of TMS-EEG, a grand-average TEP cannot be assumed to represent evoked responses that are reproducible across subjects. One reason for this is the strong dependence of TEP waveforms on subject-specific factors that are hard to control, such as the TMS target location relative to individual anatomy, the angle and direction of the induced electric field in relation to axonal orientation, and the intensity of the induced field on the cortical surface (Belardinelli et al., 2019; Casarotto et al., 2010; De Goede et al., 2018; Janssen et al., 2015; Rossini et al., 2015; Schmidt et al., 2015).

In the present study we analysed two motor-cortex TMS–EEG datasets collected using state-of-the-art practices to control for TEP quality. Both groups used adaptable noise masking to suppress auditory artifacts (Russo et al., 2022), neuronavigation to control coil position and angle (Lioumis and Rosanova, 2022), real-time TEP visualisation to monitor response quality during acquisition (Casarotto et al., 2022) and individualized procedures to set stimulation parameters. In the Milan cohort, stimulation parameters were adjusted individually on the basis of TEP amplitude in real time, whereas in the Aalborg cohort, stimulation intensity was determined using a fixed percentage of the subject’s resting motor threshold. Despite these precautions, both cohorts exhibited pronounced inter-subject variability in EEG responses, including polarity inversions across subjects (Figure 3). This is not unexpected given that the same scalp electrodes rarely sample identical neural sources across individuals (Scrivener and Reader, 2022) and anatomical differences can significantly alter induced electric field and TEP morphology (Vlachos et al., 2022; Lee et al., 2018). In such circumstances a simple grand average is unlikely to recover the underlying reproducible components (Ozdemir et al., 2021).

GTRCA addresses this problem by accommodating spatial differences across subjects, rather than averaging them away. By employing subject-specific spatial filters to identify components with maximal temporal covariance across trials both within and between subjects, gTRCA can detect a component that is reproducible even when its polarity differs between participants to the point that grand averaging cancels it entirely (Figure 2). This flexibility distinguishes gTRCA from other multivariate methods for characterizing evoked potentials, such as Correlated Component Analysis (Parra et al., 2019) and similar techniques (Zhuang et al., 2020), which impose a single spatial filter on all individuals.

In our data, polarity inversions and spatial variability made the grand-average TEPs of the two cohorts poorly comparable (Figure 4C). Yet, the first, most robust gTRCA component extracted from each cohort was reproducible across trials, across subjects, and – critically – across both datasets (Fig. 7C). Notably, this same component matched well-known signatures of M1 TEPs reported by various independent groups (Beck et al., 2024; Farzan and Bortoletto, 2022; Fecchio et al., 2017; Rocchi et al., 2021): its peak latencies (around 15ms, 30ms, 50-60ms, and 110-120ms; Figures 5C and 7A) reflected the reported early peaks of motor TEPs (Ahn and Fröhlich, 2021; Beck et al., 2024; Bonato et al., 2006; Komssi et al., 2002; Van Der Werf and Paus, 2006); its topography was centered over the stimulated area (Figures 6 and 7A) and its spectrum peaked in the low beta range, typical of primary sensorimotor cortex (Fecchio et al., 2017; Hannah et al., 2022; Van Der Werf and Paus, 2006). Together, these observations indicate that gTRCA is able to automatically extract components that most likely reflect genuine cortical responses to TMS (Belardinelli et al., 2019; Fecchio et al., 2025).

A valuable consequence of gTRCA’s flexibility as a dimensionality reduction method is its ability to simultaneously capture common group patterns and flag participants whose topographies deviate from the group norm (Figure 6). This dual sensitivity - to shared features and to meaningful outliers - is essential for developing reliable clinical biomarkers (Loth et al., 2021) and may prove especially valuable in cohorts of patients with focal or multifocal brain lesions, where the same TEP component may originate from anatomically distinct cortical regions in different individuals (Sarasso et al., 2020).

### 4.2 Quantifying the level of reproducibility of a group of TEPs with gTRCA

Each gTRCA component is paired with an eigenvalue that quantifies group reproducibility, i.e. how closely the component’s time course aligns across trials and across participants. Proper interpretation of gTRCA eigenvalues, however, requires an appropriate statistical framework. Most multivariate approaches for evoked potentials aim to optimize temporal similarity along a single dimension (trials or subjects), and statistical methods for determining the significance of their eigenvalues can thus be based on simple null hypotheses, such as the absence of time-locking across trials (Parra et al., 2019; Tanaka et al., 2013; Zhuang et al., 2020). In the case of gTRCA, which optimizes covariance both across trials and subjects, one should not assume that rejecting a null hypothesis formed by the conjunction of two factors implies the rejection of each of the individual factors that make up the hypothesis. Therefore, to disentangle these contributions, we introduced here two distinct statistical approaches: trial-based shifting and subject-based shifting. The results from simulated data showed that these tests address complementary aspects of the reproducibility in a set of evoked potentials: significance on the trial-based shifting suggests the existence of temporal locking in the dataset (either across trials or subjects, or both), while significance on the subject-based shifting specifically indicates reproducibility across subjects (Figure 2).

Quantifying group reproducibility with a single objective measure is valuable not only for evaluating statistical significance within a cohort, but also for comparing different groups, as many conditions of interest in TMS-EEG research are sources of variability in TEP waveforms. Inter-subject variability can reflect individual differences in brain connectivity and dynamics (Ozdemir et al., 2021) and arise from many state-dependent factors including attention level (Herring et al., 2015), alertness (Noreika et al., 2020), ongoing sensory inputs (Rutiku et al., 2013), spontaneous fluctuations of neuronal oscillations (Janssens and Sack, 2021), caffeine intake (Murd et al., 2010), time awake (Huber et al., 2013) and circadian regulation (Ly et al., 2016). Two practical caveats are worth noting in this context. First, because gTRCA eigenvalues scale linearly with the number of subjects, they should be normalized before being compared across different cohorts. Second, very small trial counts yield poor covariance estimates and spuriously inflated eigenvalues (see Supplementary Figure 5). For this reason, reliable comparisons of normalized eigenvalues are possible only when datasets contain similar numbers of trials.

### 4.3 Reliability of gTRCA TMS-evoked components

GTRCA resulted in topographies that were largely similar among subjects (Figure 6, Supplementary Figure 3), despite no spatial restriction imposed by the method. This fact is a strong indication of the potential of gTRCA in avoiding the risk of overfitting. Indeed, our results demonstrate that the first gTRCA component is highly robust to data variation, retaining consistent waveforms and topographic distributions across different subsets of participants, and even after reducing the sample size by approximately 50% (Figure 9). This observation supports the use of TMS-EEG in group-level investigations with a reduced number of participants (around ten), provided that TEPs are collected with sufficient number of trials and following procedures aimed at maximizing the impact of TMS on the cortex, minimizing sources of biological confounds, and ensuring within-session reproducibility of the stimulation parameters (Casarotto et al., 2022; Lioumis and Rosanova, 2022; Russo et al., 2022).

At the same time, our findings suggest that employing a larger sample size could uncover reproducible TEP components that are undetected with a smaller sample. For example, the third component on Figures 5 and 7 failed to reach significance in several pools of the leave-one-out procedure, in which one subject was removed from the group. Future studies should therefore investigate the existence of significant gTRCA components evoked by TMS that may not have been observable here due to the limited number of participants.

An important aspect of our results concerns the dependence of the gTRCA components on the number of single trials. Usually, extracting M1 TEPs with high signal-to-noise ratio requires about 100-150 noise-free trials (Hernandez-Pavon et al., 2023; Kerwin et al., 2018). However, since gTRCA preserves both dimensions (trials and subjects), the most reproducible gTRCA components could be reliably extracted even with substantially reduced number of trials (around 40, Figure 8). While these results should be interpreted with caution, as trial removal occurred only after signal preprocessing, the reliability of the method for extracting the principal gTRCA component with a reduced number of trials opens the possibility to facilitate the identification of TEP waveforms, especially in challenging recording conditions such as those encountered in clinical practice.

### 4.4 Limitations and future perspectives

As a final remark, here gTRCA was only tested on two sets of motor TEPs, which are a special type of TMS-EEG potential, given that targeting the motor cortex enables more precise control of stimulation parameters across subjects (Rossini et al., 2015). A first step in further validating the use of gTRCA in TMS-EEG research has been achieved in a recent study (Fecchio et al., 2025) that showed that the waveform and scalp map of the main gTRCA component can discriminate between motor and premotor stimulation, as well as between real and sham TMS. Looking ahead, applying gTRCA to TMS-EEG data from other, more challenging cortical regions – such as frontal and lateral areas (Farzan, 2024) – and to more diverse groups of participants will be crucial for evaluating its potential as a method for classifying TEPs under clinically relevant conditions.

## Data and Code Availability

The data that support the findings of this study are available upon reasonable request to the corresponding author. The gTRCA toolbox for evoked potentials is available in Python at https://github.com/Boutoo/gTRCA.

## Author Contributions

Conceptualization: BAC, AGC. Methodology: BAC, AGC. Investigation: BAC, MF, SR, EDM, SP, SS, DCA, MR, AGC. Data Curation: BAC, MF, EM, MR, AGC. Resources: TGN, DCA, MM, MR, AGC. Software: BAC, AGC. Formal analysis: BAC, AGC. Writing – original draft: BAC, AGC. Writing – review and editing: BAC, MF, SR, EDM, SP, SS, TGN, DCA, MM, MR, AGC. Visualization: BAC, AGC. Funding Acquisition: MF, TGN, DCA, MM, MR, AGC. Supervision: DCA, MM, MR, AGC.

## 5 Supplementary Material

### 5.1 Supplementary Methods

In this section we describe the main steps of the Python algorithm for extracting the spatial filters *w*, gTRCA components *y*_*α*_, and scalp maps *m*_*α*_ from epoched data 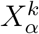 recorded using *n* EEG channels in a group of *A* individuals (*α* = 1, …, *A*) and *K* trials (*k* = 1, …, *K*). The code is available at https://github.com/Boutoo/gTRCA.

1. gTRCA is implemented in a “gTRCA class” (function gtrca.py) constructed from epochs in the MNE-Python format (Gramfort, 2013) through the gTRCA.fit() method. The first step of the procedure loads the segmented data 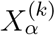, concatenates trials in the matrix *X*_*α*_ and normalizes the result, ensuring zero mean and unit variance for each channel across all subjects.
2. The second step of the gTRCA.*fit()* method calculates the matrices *S* and *Q* and proceeds with the inversion of the *Q* matrix. In the case of TMS-EEG data, pre-processing steps such as average reference, bad channels interpolation and use of ICA typically result in a low-rank covariance matrix (*rank < nA*) and strategies of regularization need to be employed. The fit method applies a Singular Value Decomposition regularization in order to detect the real dimensionality of the data before proceeding with matrix inversion.
3. In the final step, the *fit* method derives the eigenvalues (*λ*) and eigenvectors (*w*) through eigen-decomposition of the *Q*^*−*1^*S* matrix.
4. gTRCA components can then be extracted using the *get component* method of the gTRCA class. The time-series of the components are obtained for each subject through the product of the subject’s eigenvectors (spatial filters) and the epoched data. The corresponding scalp maps are calculated as the projection of the subject’s covariance matrices *Q*_*α*_ onto the eigenvectors.
5. The *get_component* method returns individual components that are both normalized and oriented. Normalization is achieved by ensuring that the average time series of each subject has unit variance and zero-mean baseline. Additionally, since the sign of eigenvectors is arbitrary, the same component can result inverted across different subjects. Proper orientation is crucial when computing average time series and scalp maps across subjects, as components with opposite orientations can cancel each other out during averaging. In the original gTRCA paper, eigenvectors were oriented by the sign of the correlation between the correspondent components and the signal at the Oz channel (Tanaka, 2020). Here we employed a two steps procedure: first, we verified that the peak of each individual component was aligned with the polarity of the corresponding peak at the group-level (i.e., the mean across subjects). Then, we recalculated the group average and orientated individual components based on the sign of its correlation with this group average. This procedure can be applied either in the temporal domain (in which components are oriented by their time courses) or in the spatial domain (in which components are oriented by their spatial maps). The *get_component* method uses temporal orientation for averaging time courses and spatial orientation for averaging spatial maps.

### 5.2 Supplementary Figures

**Supplementary Figure 1.**
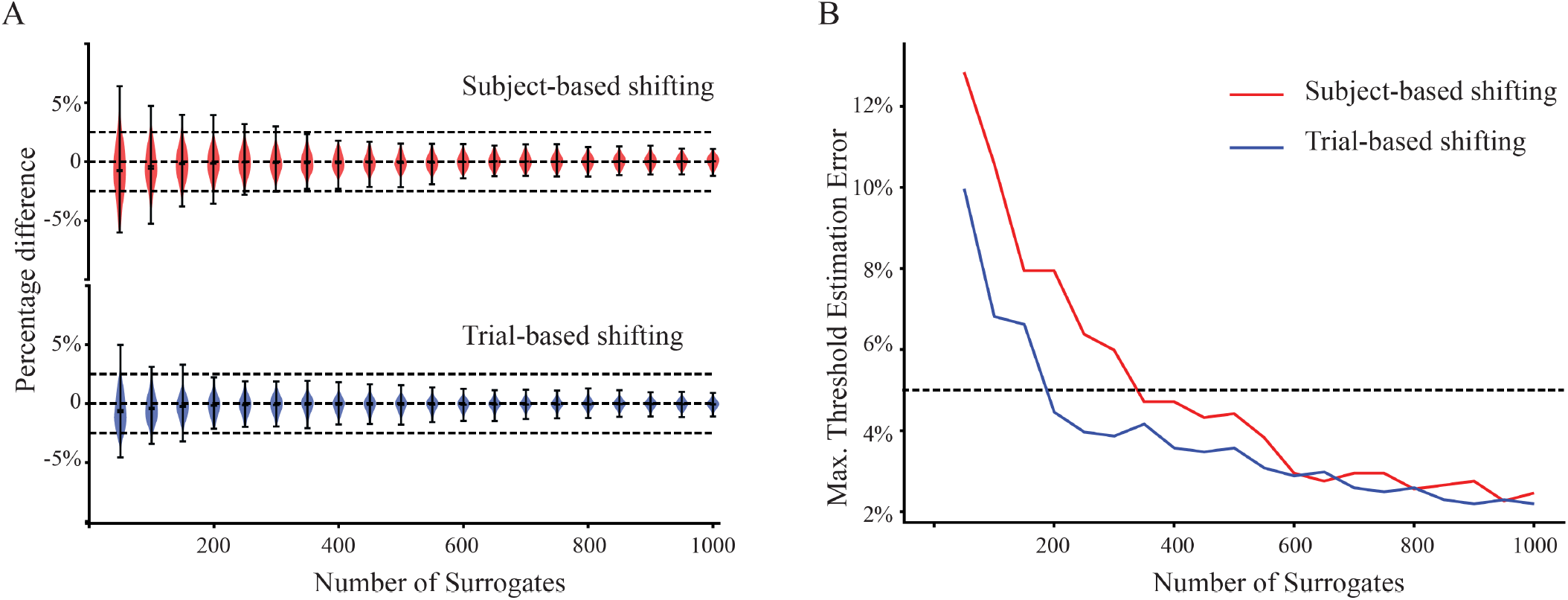
**(A)** Error in estimating the gTRCA statistical thresholds for the Milan dataset when using a reduced number of surrogates (N). For each N, the full set of 5,000 surrogates was subsampled 200 times, and a threshold was recalculated for every subsample (subject-based test in red, trial-based test in blue). Error is expressed as the percentage difference between each subsampled threshold and the reference threshold obtained with 5,000 surrogates. Violin plots depict the distribution of percentage differences; horizontal dashed lines mark ±5% error. **(B)** Maximum percentage deviation observed across the 200 subsamples for each N (red = subject-based; blue = trial-based). The dashed line indicates a 5% deviation. In the subject-based test, approximately 400 surrogates ensured less than 5% error in every iteration; in the trial-based test, around 200 surrogates were sufficient. All 4,000 surrogate-based iterations reproduced exactly the same three between-subject components reported in Figure 5 of the main text.

**Supplementary Figure 2.**
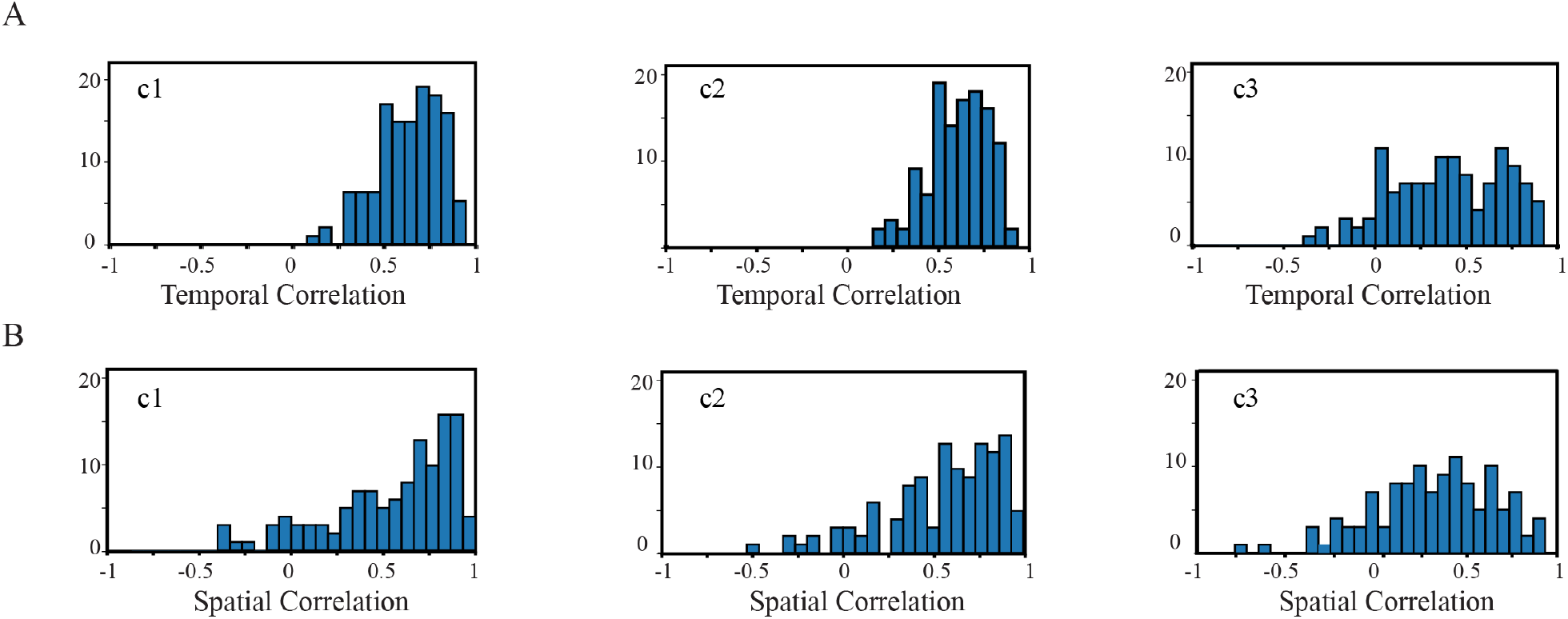
Distributions of temporal (**a**) and spatial (**B**) correlation values for each significant TEP component evoked by M1 stimulation across all subjects in the Milan dataset, taken pairwise (c1 left; c2 in the middle; c3 right). Components were oriented according to the type of correlation, whether on time (A) or space (B).

**Supplementary Figure 3.**
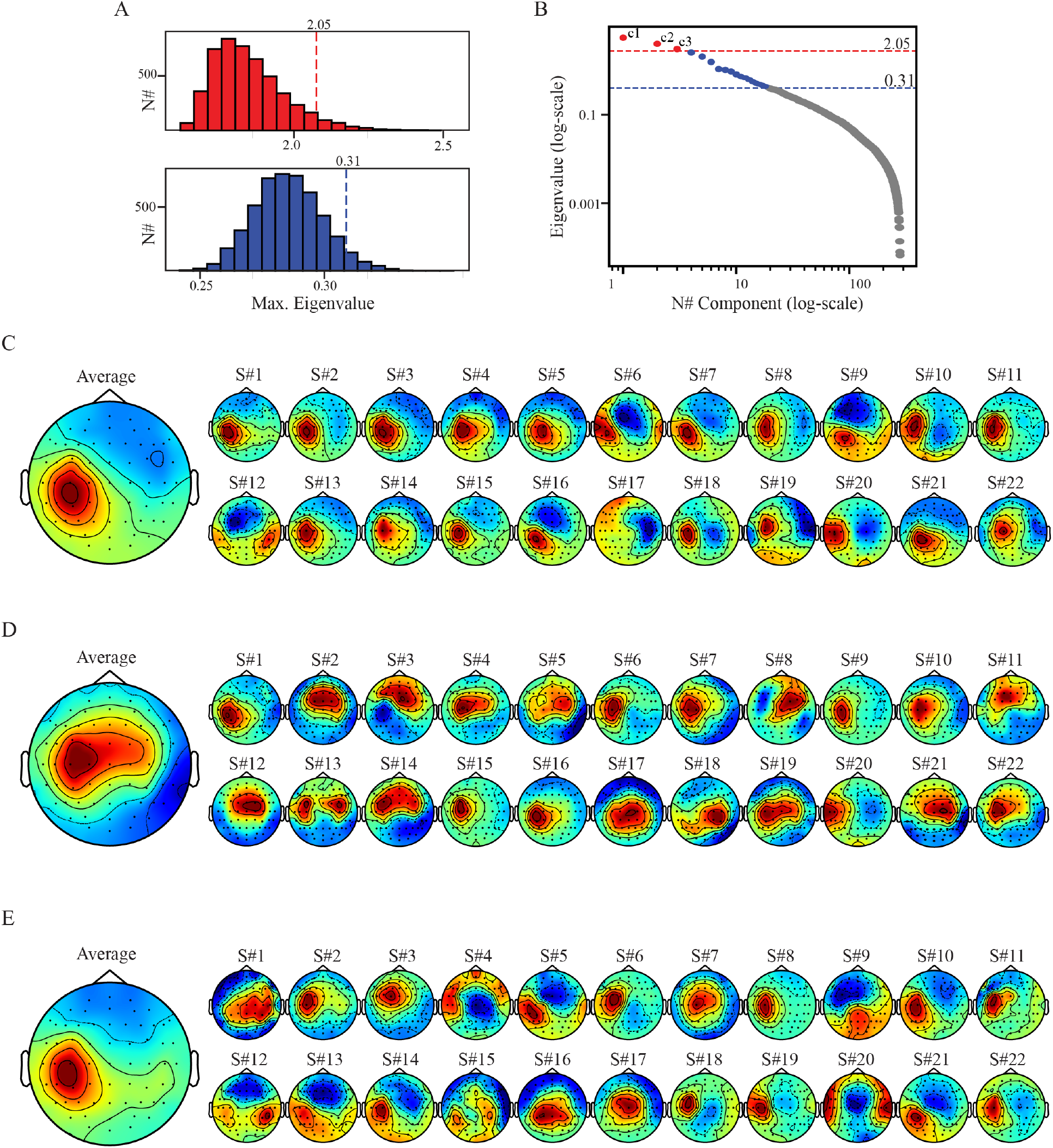
**(A)** Statistical null-distributions of gTRCA eigenvalues for each method in the Aalborg dataset: trial-based shifting (blue) and subject-based shifting (red). The dashed lines indicate the 95% quantiles, with the corresponding values of the threshold exhibited on top. **(B)** Eigenvalues (in logarithmic scale) obtained from the application of gTRCA. Components that were found not significant in both tests are colored in gray, those that were significant only in the trial-based shifting are colored in blue, and those that were significant in both tests (trial and subject-based shifting) are colored in red and marked as c1, c2 and c3. Vertical dashed lines exhibit the thresholds for the trial-based (blue) and subject-based (red) shifting methods. **(C-D-E)** Scalp maps for each significant component (C: c1; D: c2, E: c3). Averages across all subjects are shown in the left topographical plots, with the corresponding individual topographical maps displayed to the right.

**Supplementary Figure 4.**
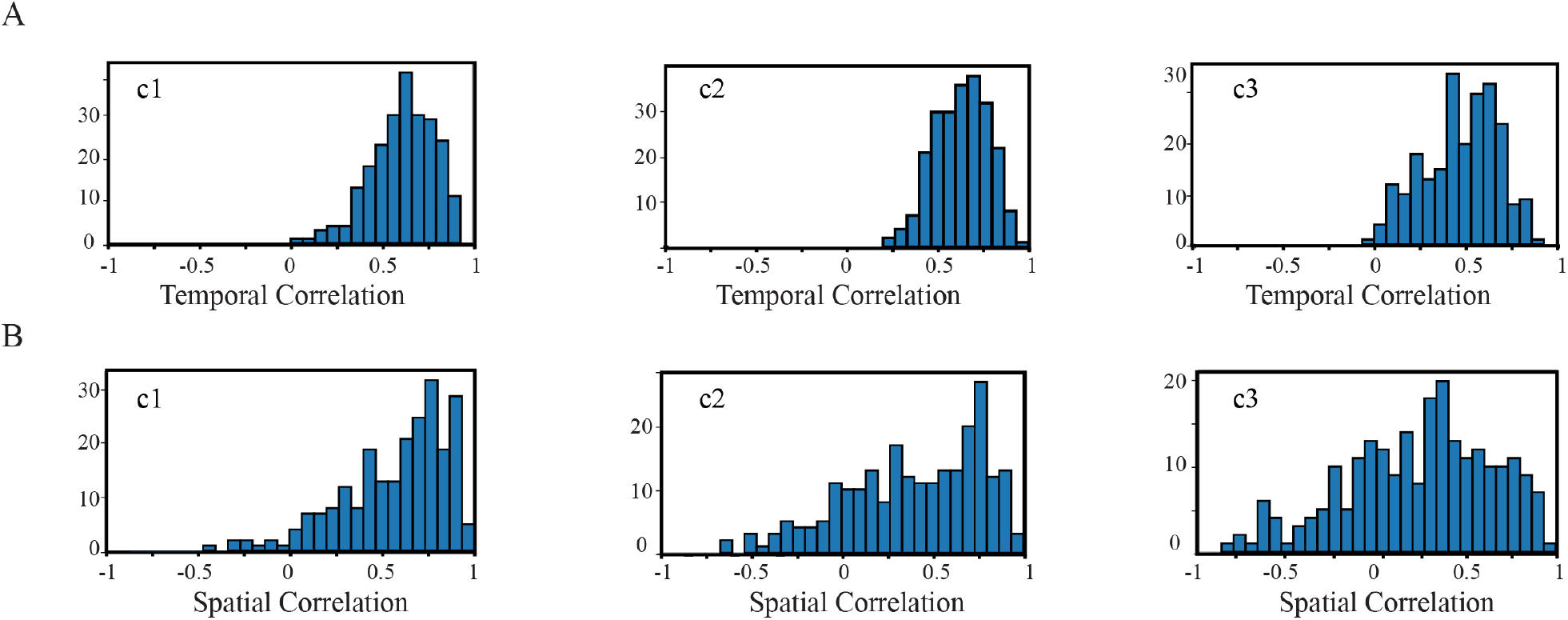
Distributions of temporal **(A)** and spatial **(B)** correlation values for each significant TEP component evoked by M1 stimulation across all subjects in the Aalborg dataset, taken pairwise (c1 left; c2 in the middle; c3 right). Components were oriented according to the type of correlation, whether on time (A) or space (B).

**Supplementary Figure 5.**
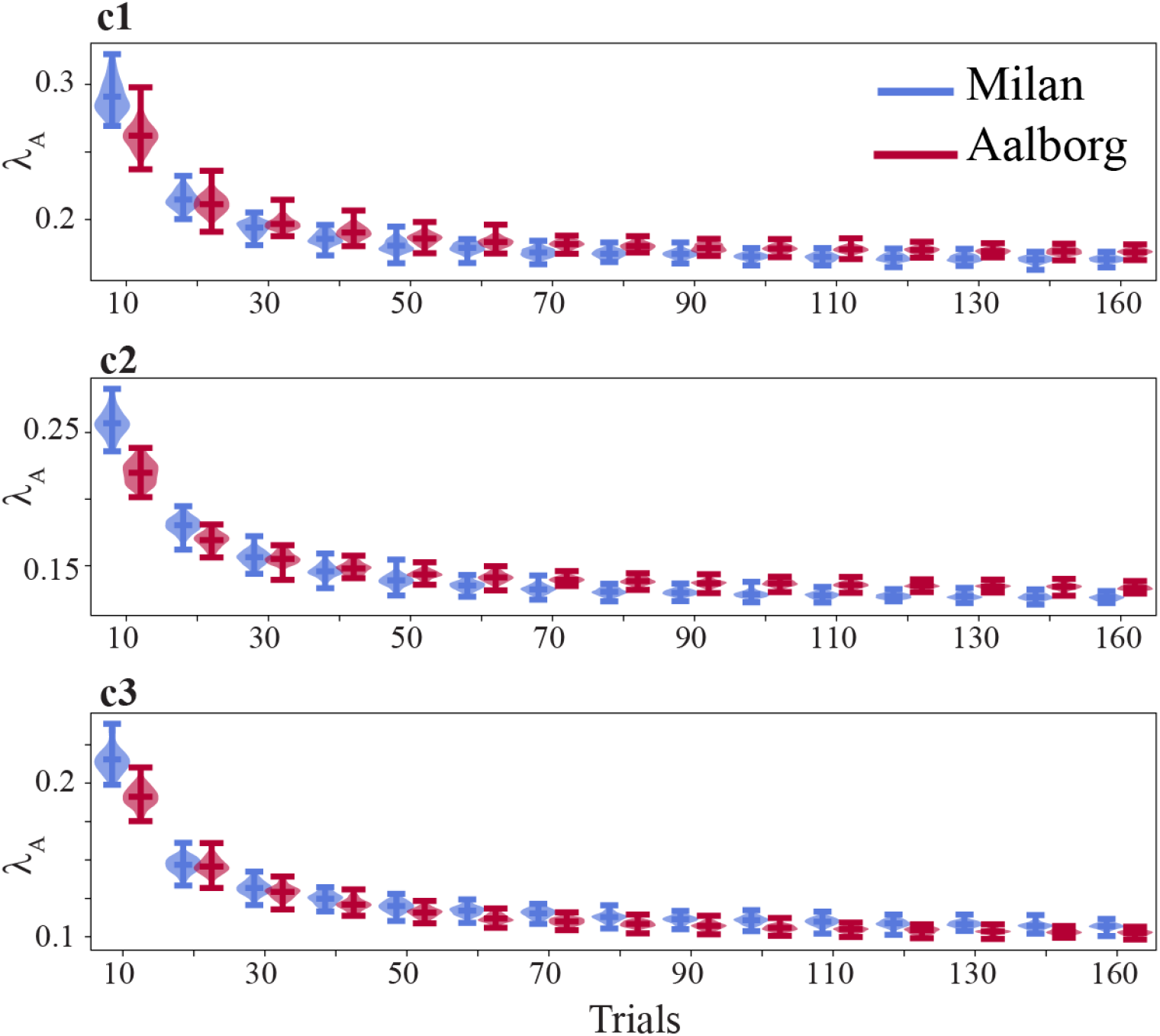
Violin plots of normalized eigenvalues (*λ*_*A*_) for the Milan (light blue) and Aalborg (dark red) datasets as a function of the number of trials. Each graph shows the mean eigenvalue, along with the 2.5th and 97.5th percentiles for each significant component (c1 above, c2 in the middle, c3 at the bottom).

